# Structural-Functional Brain Network Coupling Predicts Human Cognitive Ability

**DOI:** 10.1101/2023.02.09.527639

**Authors:** Johanna L. Popp, Jonas A. Thiele, Joshua Faskowitz, Caio Seguin, Olaf Sporns, Kirsten Hilger

## Abstract

Individual differences in general cognitive ability (GCA) have a biological basis within the structure and function of the human brain. Network neuroscience investigations revealed neural correlates of GCA in structural as well as in functional brain networks. However, whether the relationship between structural and functional networks, the structural-functional brain network coupling (SC-FC coupling), is related to individual differences in GCA remains an open question. We used data from 1030 adults of the Human Connectome Project, derived structural connectivity from diffusion weighted imaging, functional connectivity from resting-state fMRI, and assessed GCA as a latent *g*-factor from 12 cognitive tasks. Two similarity measures and six communication measures were used to model possible functional interactions arising from structural brain networks. SC-FC coupling was estimated as the degree to which these measures align with the actual functional connectivity, providing insights into different neural communication strategies. At the whole-brain level, higher GCA was associated with higher SC-FC coupling, but only when considering path transitivity as neural communication strategy. Taking region-specific variations in the SC-FC coupling strategy into account and differentiating between positive and negative associations with GCA, allows for prediction of individual cognitive ability scores in a cross-validated prediction framework (correlation between predicted and observed scores: *r* = .25, *p* < .001). The same model also predicts GCA scores in a completely independent sample (*N* = 567, *r* = .19, *p* < .001). Our results propose structural-functional brain network coupling as a neurobiological correlate of GCA and suggest brain region-specific coupling strategies as neural basis of efficient information processing predictive of cognitive ability.

## 1. Introduction

Humans differ in their level of general cognitive ability (GCA), frequently assessed by measures of intelligence. Although GCA is often interchangeably referred to as general intelligence (Kovacs & Conway, 2019; Plomin, 1999; Sternberg, 2019), GCA represents a rather vague and often poorly-defined construct of cognitive functioning, while the psychological concept of general intelligence is distinguishable from other concepts of intelligence and has a more precise theoretical foundation. Specifically, it is based on the observation that performance scores on all kinds of cognitive tasks are positively correlated with one another, i.e., there exists a positive manifold. According to the *g*-factor theory of intelligence (Spearman, 1904), the performance on each task is determined by general intelligence (a single underlying intelligence factor called the *g*-factor) reflecting the latent mental ability dimension that is common to all tasks, and by a specific factor *s* that is unique to each given task. Research across the last decades demonstrated that individual differences in general intelligence (and in GCA) are associated with important life outcomes including academic and occupational achievement (Deary et al., 2010), socio-economic status (Strenze, 2007), and even with health and longevity (Deary et al., 2004). Although intact brain structure and brain function are essential for effective cognition (Woolgar et al., 2010), the neurobiological mechanisms underlying individual differences in GCA remain elusive (Barbey et al., 2021; Basten et al., 2015; Hilger et al., 2022). Network neuroscience theories of intelligence propose that not only the structure and function of distinct brain regions, but especially the interactions and the information flow between them is critical to explain individual differences in intelligence (Barbey, 2018; Hilger and Sporns, 2021). Such conceptual models are closely related to psychological theories which postulate that GCA results from coordinated action of several fundamental cognitive processes (including, e.g., working memory capacity and mental processing speed; e.g., Duncan et al., 2020; Frischkorn et al., 2019; McKinney and Euler, 2019; for review see Hilger et al., 2022).

Support for network neuroscience theories comes from studies relating individual differences in GCA to various characteristics of structural brain network connectivity (SC; for a comprehensive overview see Genç and Fraenz, 2021) including e.g., whole-brain white-matter integrity (Chiang et al., 2009; Navas-Sánchez et al., 2014; Penke et al., 2012). Characteristics of functional brain network connectivity (FC) have also been linked to GCA such as, for example, the efficiency and the modularity of brain regions implicated in higher cognitive functions (e.g., Bertolero et al., 2018; Finn et al., 2015; Hilger et al., 2017, 2020; Kruschwitz et al., 2018; Thiele et al., 2022; for a comprehensive overview see Hilger and Sporns, 2021). However, how the alignment of the two modalities – the structural-functional brain network coupling (SC-FC coupling) – relates to GCA has not yet been investigated.

While SC and FC are significantly correlated (i.e., coupled), there is imperfect correspondence (Suárez et al., 2020). Various methods have been developed to estimate the amount of SC-FC coupling, including statistical models (Messé et al., 2014; Mišić et al., 2016), biophysical models (Breakspear, 2017; Deco et al., 2009; Honey et al., 2007), and communication models (Crofts and Higham, 2009; Goñi et al., 2014; Mišić et al., 2015). A straight-forward statistical approach is to directly compare both modalities by correlating structural and functional connectivity matrices (Baum et al., 2020; Gu et al., 2021). However, one of the main challenges to this approach is that SC represents a sparse matrix that only captures direct anatomical connections, while FC represents a full matrix that captures all pairwise interactions regardless of direct anatomical linkage. Overcoming this problem requires a model of neural dynamics that can be applied on the sparse SC matrix and approximates relationships between brain regions that are not directly structurally connected. Similarity measures are one type of such model expressing the similarity of structural connections between all possible pairs of brain regions. Their application results in almost fully connected similarity matrices thus bridging the gap between brain structure and function (Zamani Esfahlani et al., 2022). Other studies assessed SC-FC coupling indirectly by focusing on SC-behavior relationships and FC-behavior relationships separately while subsequently identifying overlapping brain connections (Dhamala et al., 2021; Zimmermann et al., 2018). Biophysical models consider plausible biological mechanisms to model the link between SC and FC but are computationally costly (Murray et al., 2018; Suárez et al., 2020). Lastly, network communication models are another way to estimate neural dynamics from SC based on specific strategies of neural communication (e.g., shortest path routing, diffusion, or navigation). Like in the case of similarity measures, this approach also results in nearly fully connected communication matrices (computed from SC) that can then be compared to the actual FC, so that both approaches allow for a valid examination of SC-FC coupling (Abdelnour et al., 2014; Seguin et al., 2020; Suárez et al., 2020).

More specifically, communication measures quantify the ease of communication between pairs of brain regions under the signaling strategy proposed by a specific communication model (Seguin et al., 2022). In contrast to statistical approaches only quantifying the amount of SC-FC coupling, the degree to which communication measures (computed on the basis of the SC) overlap with the actual FC provides insights into different neural communication processes (Avena-Koenigsberger et al., 2018; Betzel et al., 2022; Goñi et al., 2014; Rubinov and Sporns, 2010; Seguin et al., 2020, 2022; Zamani Esfahlani et al., 2022). Support for the utility of communication models to investigate SC-FC coupling comes from studies reporting improved coupling strength when communication measures, instead of the raw SC, were set in relation to FC (Goñi et al., 2014; Seguin et al., 2020, 2022) and from research examining SC-FC coupling with respect to brain development (Zamani Esfahlani et al., 2022) and human behavior (Seguin et al., 2020). Finally, communication models have also been proposed as promising means of analyzing SC-FC coupling with respect to individual differences (Avena-Koenigsberger et al., 2018; Goñi et al., 2014; Seguin et al., 2020).

The increased interest in SC-FC coupling motivated research on relationships with age (Baum et al., 2020; Hagmann et al., 2010), gender (Gu et al., 2021; Zhao et al., 2021), heritability (Gu et al., 2021), and disease (Ma et al., 2021; Rui et al., 2020; H. Zhang et al., 2021, X. Zhang et al., 2022). Also, there have been first efforts to investigate the relationship between SC-FC coupling and individual differences in cognitive ability: While stronger SC-FC coupling has been related to decreased cognitive functioning (Wang et al., 2018), other studies found increased SC-FC coupling to promote specific processes of cognitive flexibility (Medaglia et al., 2018) and complex cognition (Griffa et al., 2022). Notably, these three studies focused on whole-brain or brain-network-wise SC-FC coupling, i.e., coupling values were averaged across the whole brain or across large brain networks (Yeo et al., 2011). In contrast, Baum et al. (2020) demonstrated that the association between SC-FC coupling and cognitive ability differs critically between brain regions, e.g., higher executive functioning was associated with increased alignment in the rostrolateral prefrontal cortex, posterior cingulate and medial occipital cortex but with decreased alignment in the somatosensory cortex. However, insights of their study are limited due to the purely statistical approach of directly correlating SC and FC to assess their coupling and by restricting analyses to only one very specific cognitive ability measure. Whether a) whole-brain SC-FC coupling is related to general cognitive ability, b) potential associations are positive or negative, and c) whether potential associations differ between distinct brain regions has not yet been investigated.

Here, we systematically examine the association between GCA and SC-FC coupling in a sample of 1030 adults from the Human Connectome Project (HCP, Van Essen et al., 2013). GCA was estimated as a latent *g*-factor derived from 12 cognitive performance measures and SC-FC coupling was operationalized with two similarity measures and six communication measures. First, we tested for potential associations between GCA and SC-FC coupling on a brain-average level. Second, a cross-validated prediction framework was developed that accounts for region-specific variations in coupling strategies as well as for positive and negative associations with GCA. This model was evaluated for its ability to predict individual cognitive ability scores in previously unseen participants. All analyses were finally repeated in an independent replication sample and the generalizability of the prediction model was assessed with a cross-sample model generalization test.

## 2. Methods

### 2.1. Preregistration

Analysis plans and variables of interest were preregistered in the Open Science Framework: https://osf.io/wr9aj. Please note that in deviation to our preregistration, the HCP was used as main sample as the initial sample did not contain all data required for the planned analyses and the benefit of having more than 1000 subjects (as contained in the HCP) was essential for developing a cross-validated prediction framework, the latter of which increases the robustness of results and allows to estimate the generalizability of our findings but was not initially planned. Also, an additional sample for external replication (AOMIC) was included. Note further, that in order to keep a clear focus, we also deviated from our preregistration in exclusively reporting the result of the first proposed hypothesis (H1), while the other stated hypotheses (H2 – H5) focusing on additional cognitive measures and potential mediating factors will be addressed in separate publications.

### 2.2. Participants

Main analyses were conducted in the HCP Young Adult Sample S1200 (details see Van Essen et al., 2013) including 1200 subjects of age 22-37 (656 female, 1089 right-handed, mean age = 28.8 years).

Subjects with missing resting-state fMRI data (from all four scans), missing DWI data, missing cognitive measures required to calculate a latent general cognitive ability factor, or a Mini-Mental State Examination (MSSE) score equal to or smaller than 26 were excluded. Further, subjects were ruled out based on in-scanner head motion measured by framewise displacement (Jenkinson et al., 2002). Following Parkes et al. (2018), scans with a) a mean framewise displacement above 0.2 mm, b) a proportion of motion spikes (framewise displacement > 0.25 mm) greater than 20 percent, or c) any spikes above 5 mm were removed. The resulting sample referred to as main sample consisted of 1030 subjects (age range 22-37, 555 female, 935 right-handed, mean age = 28.7 years).

### 2.3. General cognitive ability (GCA)

GCA was operationalized as latent *g*-factor derived from 12 cognitive measures (Table 1, see Thiele et al., 2022). The *g*-factor was calculated as outlined in Dubois et al. (2018) using simplified bi-factor analysis based on the Schmid-Leiman transformation (Schmid and Leiman, 1957).

**Table 1.**
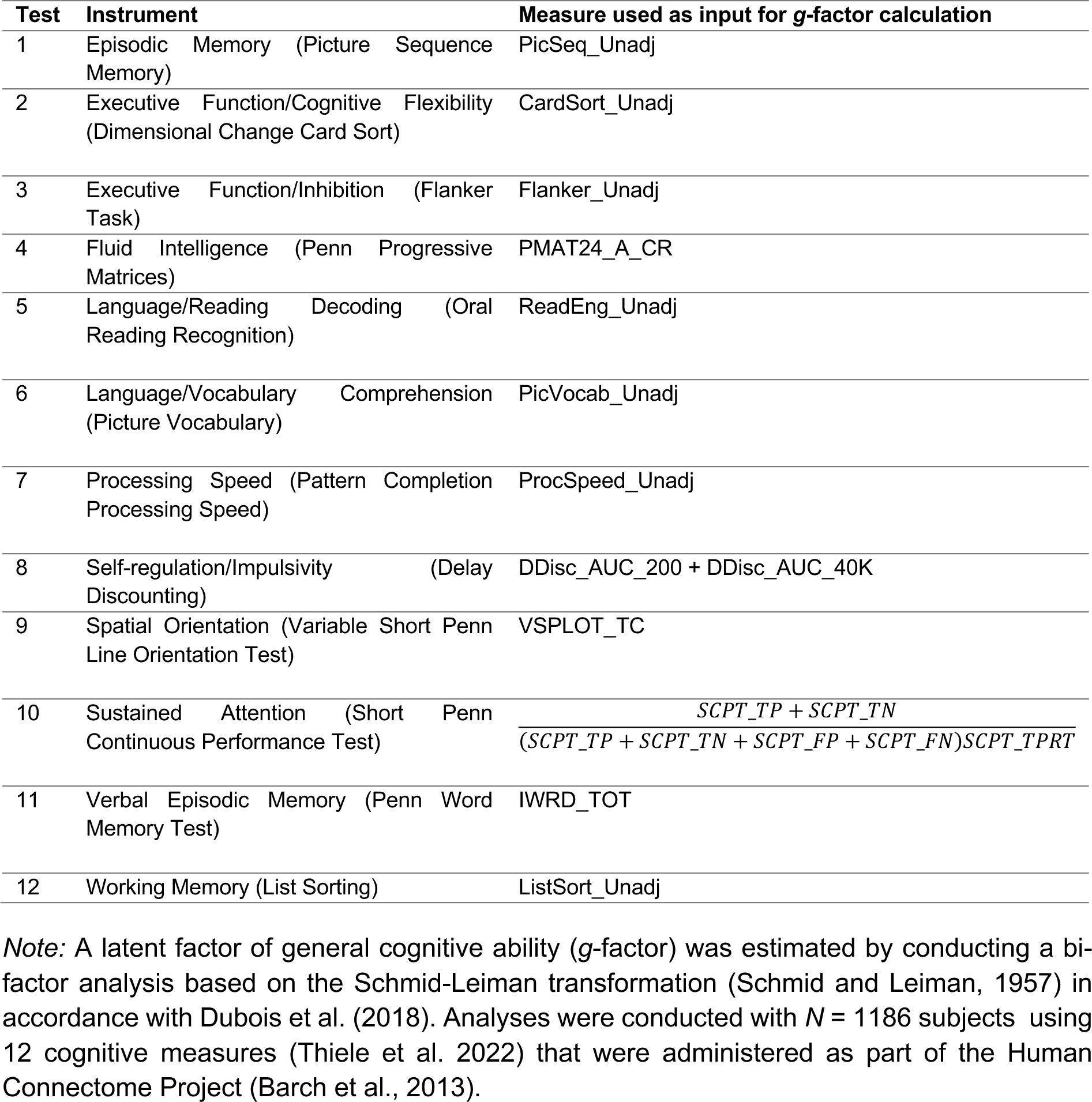
Cognitive tests and measures used to calculate a latent *g*-factor as estimate of GCA.

### 2.4. Data acquisition and preprocessing

MRI data were acquired with a gradient-echo EPI sequence on a Siemens Skyra 3T scanner with a 32-channel head coil. The fMRI scans were obtained with multi-slice acceleration (time to repetition (TR) = 720 ms, time to echo (TE) = 33.1 ms, 2-mm isotropic voxel resolution, flip angle = 52°, and multiband acceleration factor = 8). For functional brain network connectivity (FC) estimation, the minimally preprocessed resting-state fMRI data from the HCP (Glasser et al., 2013) were used. As additional denoising strategy, nuisance regression as explained in Parkes et al. (2018, strategy no. 6) with 24 head motion parameters, eight mean signals from white matter and cerebrospinal fluid and four global signals was applied. To estimate structural brain network connectivity (SC), we used data from the minimally preprocessed DWI (TR = 5520 ms, TE = 89.5 ms, 1.25 mm isotropic voxel resolution, multiband acceleration factor = 3, b = 1000, 2000, 3000 s/mm^2^, and 90 directions/shell) provided by the HCP (Glasser et al., 2013; Van Essen et al., 2013) and ran the MRtrix pipeline for DWI processing (Civier et al., 2019; Tournier et al., 2019), which includes bias correction, modeling of white matter fibers via constrained spherical deconvolution (Tournier et al., 2007), and tissue normalization (Dhollander et al., 2021). Probabilistic streamline tractography was carried out to render streamlines through white matter which terminate in grey matter (R.E. Smith et al., 2012). Additionally, filtering of streamlines was performed to only retain the streamlines that fit the estimated white matter orientations from the diffusion image (Glasser et al., 2013; R. E. Smith et al., 2013; Tournier et al., 2012).

### 2.5. Functional and structural brain network connectivity

Functional and structural brain networks were constructed by first dividing the brain into 360 cortical regions based on the multimodal parcellation scheme of Glasser et al. (2016). Note that two brain regions (left and right hippocampus) were excluded as they were regarded as subcortical regions in the preprocessing pipeline, thus resulting in 358 regions (i.e., nodes). Individual-specific FC matrices were computed by using the Fisher z-transformed Pearson correlations between BOLD time courses extracted from all possible pairs of brain regions. FC matrices were first constructed for all available resting-state scans separately and averaged afterwards (Cole et al., 2014; S. M. Smith et al., 2013). SC matrices were symmetric and defined by the SIFT2 streamline density weights between all pairs of brain regions (R.E. Smith et al., 2015).

### 2.6. SC-FC coupling

SC-FC coupling was operationalized by comparing each individual FC matrix with eight matrices. These eight matrices were computed based on the individual SC matrix using two major approaches to model potential functional interactions arising from SC, i.e., similarity and communication models. Table 2 provides a short description of all resulting similarity and communication measures.

**Table 2.**
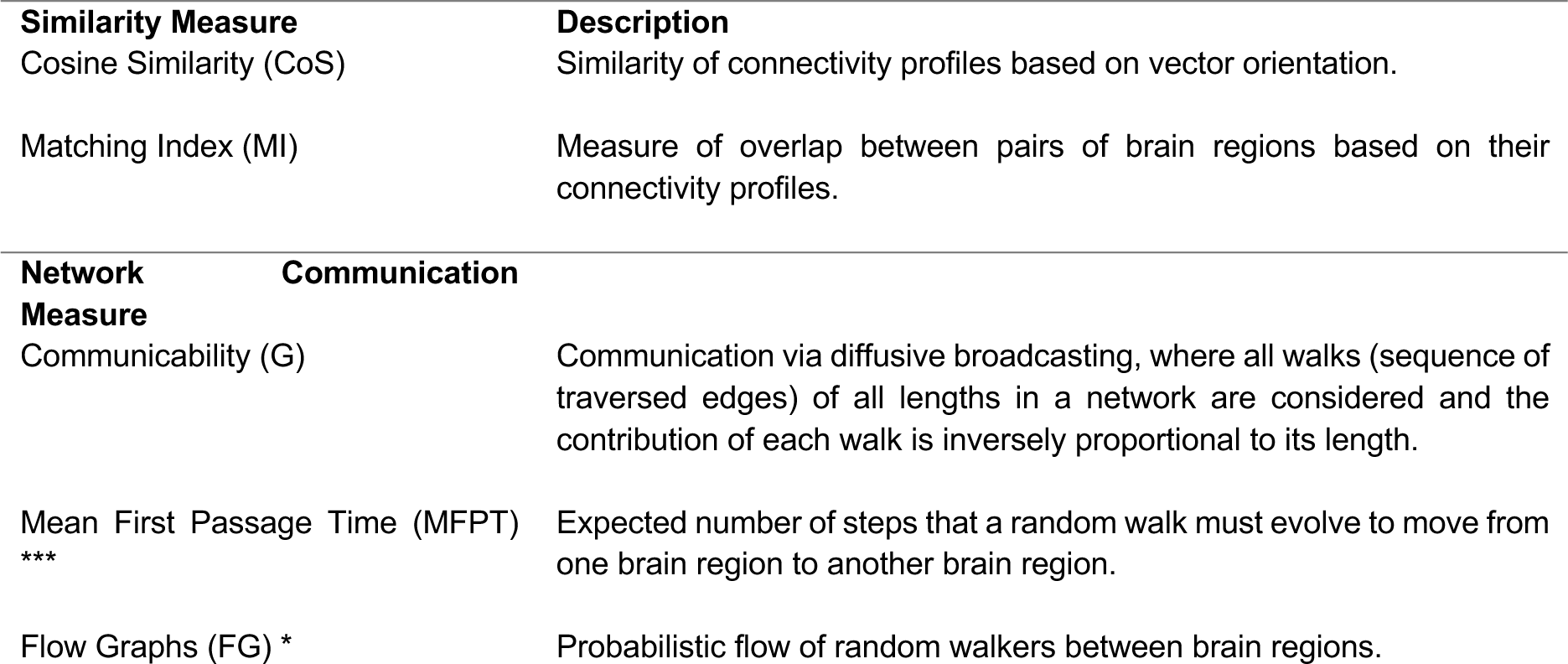
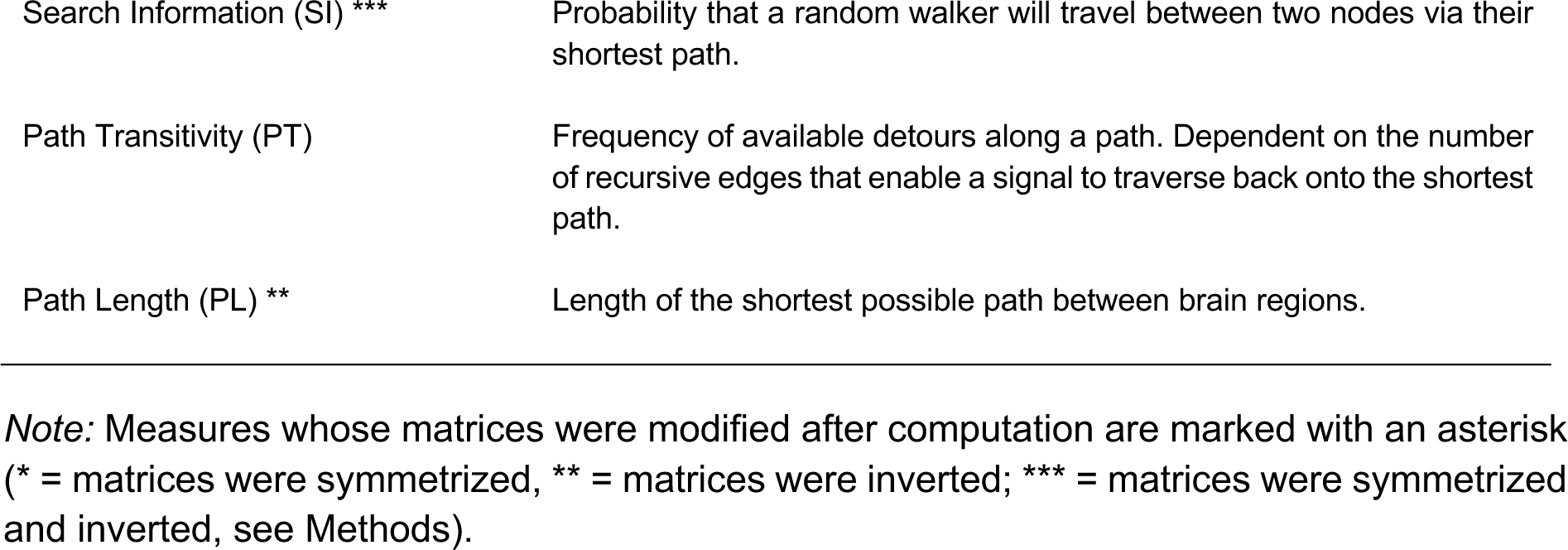
Overview of the two similarity measures and the six communication measures used to operationalize structural-functional brain network coupling.

#### 2.6.1. Similarity measures

In general, similarity measures are computed based on the SC matrix and represented in similarity matrices that express the resemblance of regional structural connectivity profiles. More specifically, an entry in the similarity matrix reflects how the structural connections of brain region *i* (defined by a matrix column) align with the structural connections of brain region *j* (defined by another matrix column). No additional information about putative signaling strategies is implemented in the calculation of similarity matrices. Each individual weighted SC matrix was transformed into two similarity matrices representing two distinct similarity measures. These were:

##### 2.6.1.1. Cosine similarity (CoS)

Cosine similarity assesses the resemblance between two brain regions’ connectivity profiles (matrix columns) based on their orientation in an *N* − 1 dimensional connectivity space, where *N* is the number of brain regions, i.e., 358. We computed the cosine similarity of the angle between two vectors *x* = [*x*_1_, …, *x_N_*] and *y* = [*y*_1_, …, *y_N_*] as 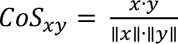, where vectors are region-specific connectivity profiles for all possible pairs of brain regions (Han et al., 2012).

##### 2.6.1.2. Matching index (MI)

Matching index measures the similarity of regional connectivity profiles between pairs of brain regions while excluding their mutual connections (Hilgetag et al., 2000, Goñi et al., 2014). For each individual SC matrix *A* with elements (matrix entries) *A_ij_*, Γ*_i_* = *j*: *A_ij_* > 0 describes the set of regions that are all directly connected to region *i*. The matching index between the two regions *i* and *j* is then calculated as 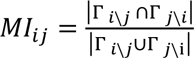, where the term Γ*_i\j_* refers to the neighbors of region *i* except region *j*.

#### 2.6.2. Communication measures

Communication measures quantify the ease of communication between pairs of brain regions under a certain signaling strategy (i.e., communication model like shortest path routing, diffusion, or navigation) and are represented in communication matrices. Specifically, each individual weighted SC matrix was transformed into six communication matrices representing six distinct communication measures. These were:

##### 2.6.2.1. Communicability (G)

Communicability considers that neural signaling unfolds as a diffusive broadcasting process, assuming that information can flow along all possible walks between two brain regions (Andreotti et al., 2014; Seguin et al., 2020). It can be defined as the weighted sum of all walks of all lengths between two respective regions (Estrada and Hatano, 2008), where an edge is the connection between two brain regions and a walk is a sequence of traversed edges. This measure accounts for all possible connections between regions but incorporates walk lengths (*l_w_*) and penalizes the contribution of walks with increased lengths. For weighted networks, SC matrices (*A*) are first normalized as *A*^’^ = *D*^-1/2^*AD*^-1/2^, where *D* is the degree diagonal matrix (Crofts and Higham, 2009). The normalized matrix is then exponentiated to calculate the communicability as *G* = *e*^A’^ or 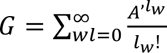, where each walk is inversely proportional to its length thus 1-step walks contribute 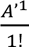, 2-step walks 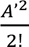 and so on.

##### 2.6.2.2. Mean first passage time (MFPT)

Mean first passage time between brain region (node) *i* and *j* refers to the expected number of steps that it takes for a random walk starting at node *i* to arrive at node *j* for the first time (Goñi et al., 2013; Noh and Rieger, 2004). If the graph of the structural brain network is defined as *G*_SC_ and composed by a set of *n* nodes *U* = {1, …, *n*}, then the graph’s connectivity is described by a *n* × *n* symmetrical connectivity matrix *A* = [*A_ij_*], where *A_ij_* defines the edges of the network and *k_i_* depicts the number of direct neighbors (*k_i_* = ∑*_j_ A_ij_*) (Goñi et al., 2013). The mean first passage time depends on a specific stochastic model, a Markov Chain, which is describing a sequence of possible events in which the probability of each event depends only on the state of the previous event (Gagniuc, 2017). A Markov Chain *M* ≡ (*St*, *Q*) consists of a set of states *St* = {*st*_1_, …, *st_n_*} and a matrix of transition probabilities *Q* = [*q_ij_*] characterizing the probability of going from one state *st_i_* to another state *st_j_* in one step (Goñi et al., 2013). A graph (i.e., structural brain network) can be expressed as a Markov chain, where states *St* = {*st*_1_, …, *st_n_*} correspond elementwise to the set of nodes *U* = {1, …, *n*}. The probability of going from one state *st_i_* to another state *st_j_* is denoted by 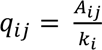, where it is assumed that there is equal probability of choosing one of the *k_i_* edges (Goñi et al., 2013). Ultimately, the mean first passage time of *G_SC_*, where nodes *U* = {1, …, *n*} stand for states *St* = {*st*_1_, …, *st_N_*} of the Markov Chain is denoted by *MFPT*_G_ = [*mfpt_ij_*] and can be computed from the fundamental matrix *Z* = [*ζ_ij_*] and the fixed row probability vector *v* as 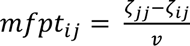, *i* ≠ *j* (Goñi et al., 2013). The probability vector *v* is the left eigenvector associated with the eigenvalue of 1 that corresponds to the stationary solution of the Markov process and the fundamental matrix *Z* is defined as *Z* = (*I* − *Q* + *V*)^-1^, where *I* is an *n* × *n* identity matrix (each element on principal diagonal is 1 and the other elements 0), *Q* is the transition matrix and *V* is an *n* × *n* matrix, where each column corresponds to the probability eigenvector *v* (Goñi et al., 2013).

##### 2.6.2.3. Flow graphs (FG)

Flow graphs are transformations of a network’s SC matrix (*A*) in which dynamic flows are embedded into the weights of edges (Lambiotte et al., 2011). More specifically, a flow graph characterizes the probability that a random walker is located at a specific position in the network between region *i* and *j* at a specific time point *t*_x_ (*Markov time* = 10 in our case). For a random walk with dynamics specified by *p_i_* = − ∑*_j_L_ij_p_j_*, a flow graph is given by *FG*(*t_m_*)*_ij_* = (*e*-*^t_m_L^*)*_i_jk_j_*.The matrix *L* is the normalized Laplacian whose elements are given by *L_ij_* = *D* − *A*/*k*, where *k* = ∑*_j_ A_ij_* is the degree of a node and *D* is the degree diagonal matrix (square matrix with a diagonal containing the elements of *k*). Thus, the variable *p_i_* represents the probability of finding a random walker on the edge between brain region *i* and brain region *j* and the element *FG*(*t_m_*)*_ij_* ultimately represents the probabilistic flow of random walkers at time *t_m_* between two respective nodes (Zamani Esfahlani et al., 2022).

##### 2.6.2.4. Search information (SI)

Search information is a measure of network navigability in the absence of global knowledge (Goñi et al., 2014) and is related to the probability that a random walker will travel between two nodes via their shortest path (i.e., the path connecting two nodes via fewest intermediate stations/nodes). This probability increases with an expanding number of paths that are available for a certain communication process to take place (Avena-Koenigsberger et al., 2018; Goñi et al., 2014). Given the shortest path between brain regions *s* (source node) and *t* (target node): *π_s_*_→*t*_ = {*s*, *i*, *j*, …, *l*, *m*, *t*}, the probability of taking this shortest path is expressed as *F*(*π_s_*_→*t*_) = *f_si_* × *f_ij_* × … × *f_lm_* × *f_mt_*, where 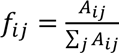 and *i*, *j*, *l* and *m* are nodes along the shortest path. The information that is then necessary to access the shortest path from *s* to *t* is *SI*(*π_s_*_→*t*_) = log_2_ [*F*(*π_s_*_→*t*_)] (Goñi et al., 2014).

##### 2.6.2.5. Path transitivity (PT)

Path transitivity also captures the accessibility of shortest paths (from source node *s* to target node *t*) within a network but accounts particularly for the frequency of detours that are available along a shortest path which would enable the signal to traverse back onto that path after leaving it (Avena-Koenigsberger et al., 2018; Goñi et al., 2014). Path transitivity is independent of the directionality of the path and defined as *PT*(*π_s_*_→*t*_) = 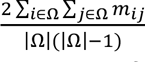, where *m_ij_* is the matching index between respective nodes on the shortest path as defined in Goñi et al. (2014).

##### 2.6.2.6. Path length (PL)

Path length is an indicator of how easily signals can be transmitted between two regions via their shortest path, since longer paths are more susceptible to noise, have longer delays in transmission and are energetically more expensive (Avena-Koenigsberger et al., 2018; Rubinov and Sporns, 2010). In a network, each edge is associated with a cost *C* (difficulty of traversing) and for weighted networks, this cost can be obtained by transforming the weight *ω* of each edge into a measure of length through *C* = *ω*^-1^. The shortest path between two respective nodes (source node *s* and target node *t)* is the sequence of edges *π_s→t_*= {*A_si_*, *A_ij_*, …, *A_mt_*} minimizing the sum *C_si_* + *C_ij_* + ⋯ + *C_mt_* (where *C_st_* is the cost of traversing the edge between region *s* and *i*) and *i*, *j* and *m* are nodes along the shortest path.

Note that as mean first passage time, path length, and search information capture difficulty of communication (instead of ease of communication like the other five measures), the respective communication matrices were transformed to ensure that each matrix entry reflects the ease of communication between brain regions and effect sizes of all measures can be interpreted in equal directions. For mean first passage time and path length, this transformation was performed by replacing all matrix entries with their element-wise reciprocal (*M_new_* = 1/*M_original_*). For SI, matrices were inverted by flipping signs for all matrix entries (*M_new_* = −1 ∗ 1/*M_original_*). The communication matrices for mean first passage time, flow graphs, and search information are asymmetric, implying that ease of communication between region *i* and *j* is not necessarily equal to the ease of communication between region *j* and *i* (Seguin et al., 2019). Thus, respective communication matrices were symmetrized to ensure that correlating matrix columns vs. matrix rows would not yield different results with regards to the regional coupling values.

For the computation of similarity and communication matrices we followed examples provided by Zamani Esfahlani et al. (2022), applying functions from the Brain Connectivity Toolbox (Rubinov and Sporns, 2010). To compute subject- and brain region-specific SC-FC coupling values, we separately compared each individual’s similarity and communication matrices to their FC matrix (one at a time; eight comparisons in total). This was done by correlating (Pearson correlation) all regional connectivity profiles (matrix columns representing the connections of one brain region to all other brain regions) of the respective similarity or communication matrix with the corresponding regional connectivity profile of the FC matrix. Each of the eight comparisons resulted in 358 individual coupling values (*r_C_*, one per brain region), thus yielding eight distinct measures approximating SC-FC coupling that are referred to as coupling measures. A workflow of the procedure is illustrated in Fig. 1.

**Fig. 1.**
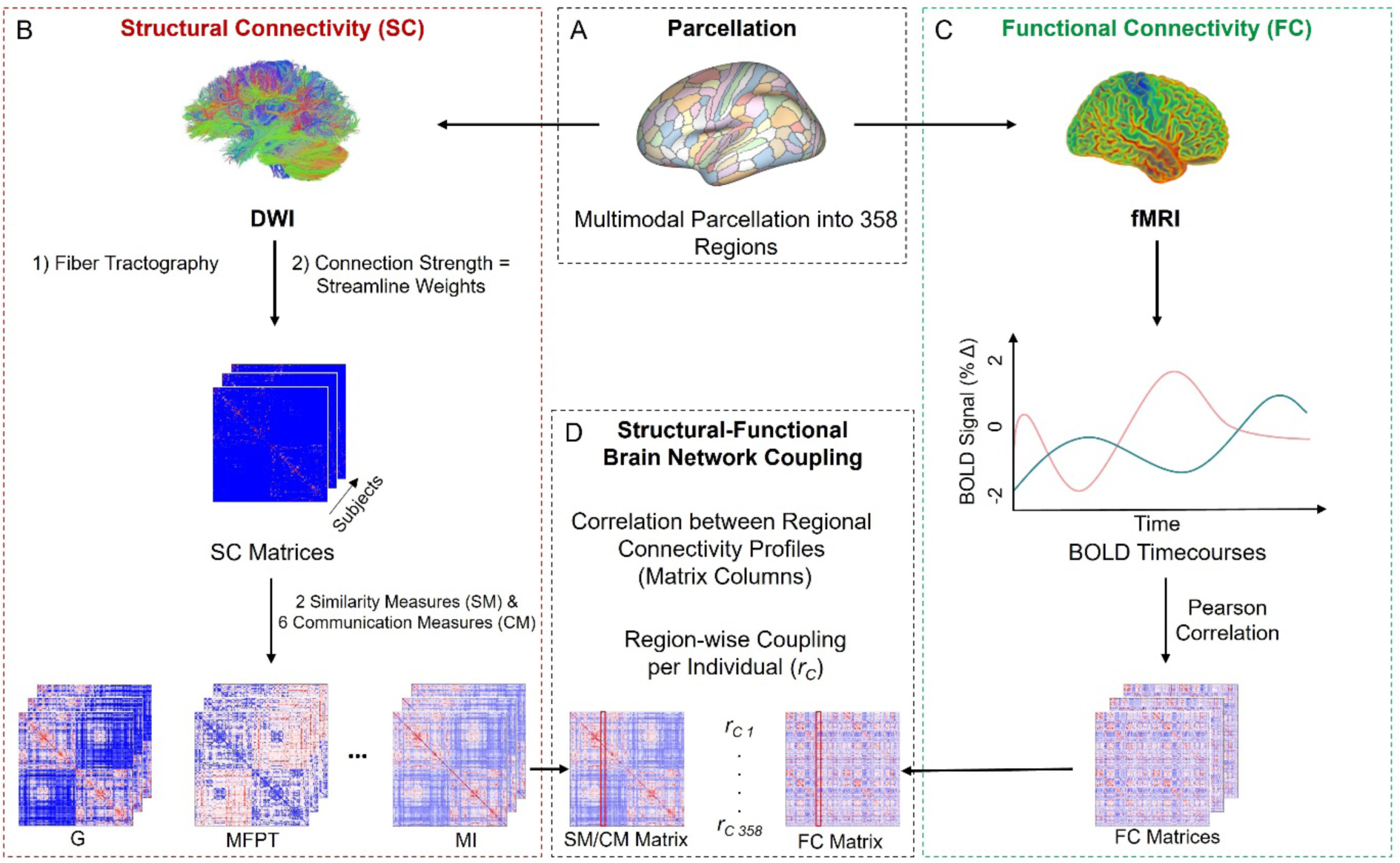
Workflow of deriving eight different measures of SC-FC brain network coupling. (A) For the operationalization of SC-FC coupling, DWI and fMRI data were parcellated into 358 brain regions based on a multimodal parcellation scheme (Glasser et al., 2016). (B) Structural connectivity matrices were transformed into similarity matrices (SM) and communication matrices (CM) expressing two distinct similarity measures and six distinct communication measures that model plausible functional interactions on top of structural connections (Table 2). (C) Functional connectivity matrices were constructed by computing Pearson correlations of regional BOLD time courses from four 15-min resting-state fMRI sessions. (D) Each individual’s two similarity matrices and each individual’s six communication matrices were then compared separately (one at a time) with the individual’s functional connectivity matrix by correlating regional connectivity profiles (matrix columns) of the respective similarity or communication matrix with the same region’s connectivity profile in the functional connectivity matrix. Each of the eight comparisons resulted in 358 individual coupling values (*r_C_*, one per brain region) thus yielding eight coupling measures. SC = Structural Brain Network Connectivity; FC = Functional Brain Network Connectivity; BOLD = Blood Oxygen Level Dependent; SM = Similarity Measure; CM = Communication Measure; G = Communicability; MFPT = Mean First Passage Time; MI = Matching Index. For illustration purposes, only three out of the eight similarity and communication matrices are shown in Fig. 1B. [color; 2-column fitting image]

### 2.7. Grouping of coupling measures

To better understand the underlying signal transmission processes, the six coupling measures derived from the comparison of communication matrices to FC matrices were grouped based on conceptual similarity of the specific signal transmission strategy that is proposed by each underlying communication model (Avena-Koenigsberger et al., 2018). Specifically, different models of neural communication can be placed on a spectrum depending on how much information is necessary for each communication process to take place. On one end of the spectrum are routing processes requiring full knowledge of network topology (e.g., target region and location of shortest paths), and on the other end are diffusive processes that operate solely on the basis of local properties (Avena-Koenigsberger et al., 2018, 2019). Thus, coupling measures were grouped into a) diffusion-based coupling measures (communicability, mean first passage time and flow graphs), b) coupling measures based on path accessibility (search information, path transitivity), and c) routing-based coupling measures (path length). Similarity-based coupling measures (cosine similarity and matching index) were considered as separate group.

### 2.8. SC-FC coupling strength across the cortex

The pattern of SC-FC coupling strength across the cortex was analyzed by first identifying the similarity or communication measure per brain region that was able to most frequently explain the highest variance (*R^2^*) in FC across all participants. This group-general mask was subsequently used to extract individual brain region-specific coupling values that were then averaged across all participants to visualize the overall SC-FC coupling pattern. Note that the approach of selecting region-specific coupling measures based on highest explained variance in FC was solely used for visualization purposes but not for further statistical analyses.

### 2.9. Brain-average SC-FC coupling

Individual coupling measure-specific brain-average coupling values were computed by taking the mean of all 358 region-specific individual coupling values (*r_C_*) for each of the eight coupling measures, previously computed by correlating region-specific connectivity profiles (as explained above). Consequently, we obtained eight brain-average coupling values per participant, one for each coupling measure.

### 2.10. Association between brain-average SC-FC coupling and GCA

To assess the relationship between brain-average SC-FC coupling and individual differences in GCA, partial correlations between cognitive ability scores and the individual brain-average coupling values (eight values per participant) were computed controlling for the influences of age, gender, handedness, and in-scanner head motion (operationalized as mean framewise displacement). Statistical significance was accepted at *p* < .05 and we corrected for multiple comparisons by applying the Bonferroni correction (eight comparisons: significant *p* < .006).

### 2.11. Association between region-specific SC-FC coupling and GCA

As previous reports revealed that the preferred communication strategy (measures best explaining FC) varies critically between different brain regions (Betzel et al., 2022; Zamani Esfahlani et al., 2022), we next developed an approach to investigate the association between SC-FC coupling and GCA on a brain region-specific level by taking this variation into consideration: For each coupling measure separately, region-specific coupling values (*r_C_*) from all participants (*N* = 1030) were correlated with individual cognitive ability scores, resulting in eight correlation coefficients (*r_G_*) per brain region reflecting brain region-specific associations of the eight coupling measures with GCA. A main goal of this approach was to examine which coupling measure per brain region was most or least able to account for individual differences in GCA. Again, partial correlations were computed controlling for age, gender, handedness, and in-scanner head motion (operationalized as mean framewise displacement).

As such correlative approaches result in extremely large numbers of comparisons (multiple comparisons problem) and are prone to overfitting (see Cwiek et al., 2022; Yarkoni and Westfall, 2017), a cross-validated predictive modeling approach was developed that a) accounts for brain region-specific differences in the preferred communication strategy and b) best prevents overfitting by creating a small number of features (instead of using all possible predictors as e.g., in a multivariate regression without regularization or feature selection) and by implementing a thorough cross-validation scheme.

#### 2.11.1. Feature construction - node-measure assignment (NMA)

The features of our prediction model were individual- and brain region-specific coupling values (*r_C_*) that were selected on the basis of group-average node-measure assignment (NMA) masks. Such group masks reflect general region-specific preferences of coupling strategies with respect to GCA. More specifically, based on the correlation coefficients (*r_G_*) from the analysis of region-specific SC-FC coupling and GCA for each brain region (see paragraph above), the coupling measures with the largest positive and negative magnitude associations with GCA were selected. This resulted in one group-based positive NMA mask and one group-based negative NMA mask, each assigning one coupling measure to one brain region and thus determining which (out of eight) individual coupling values served as input for the prediction model for a given brain region (Fig. 2A).

**Fig. 2.**
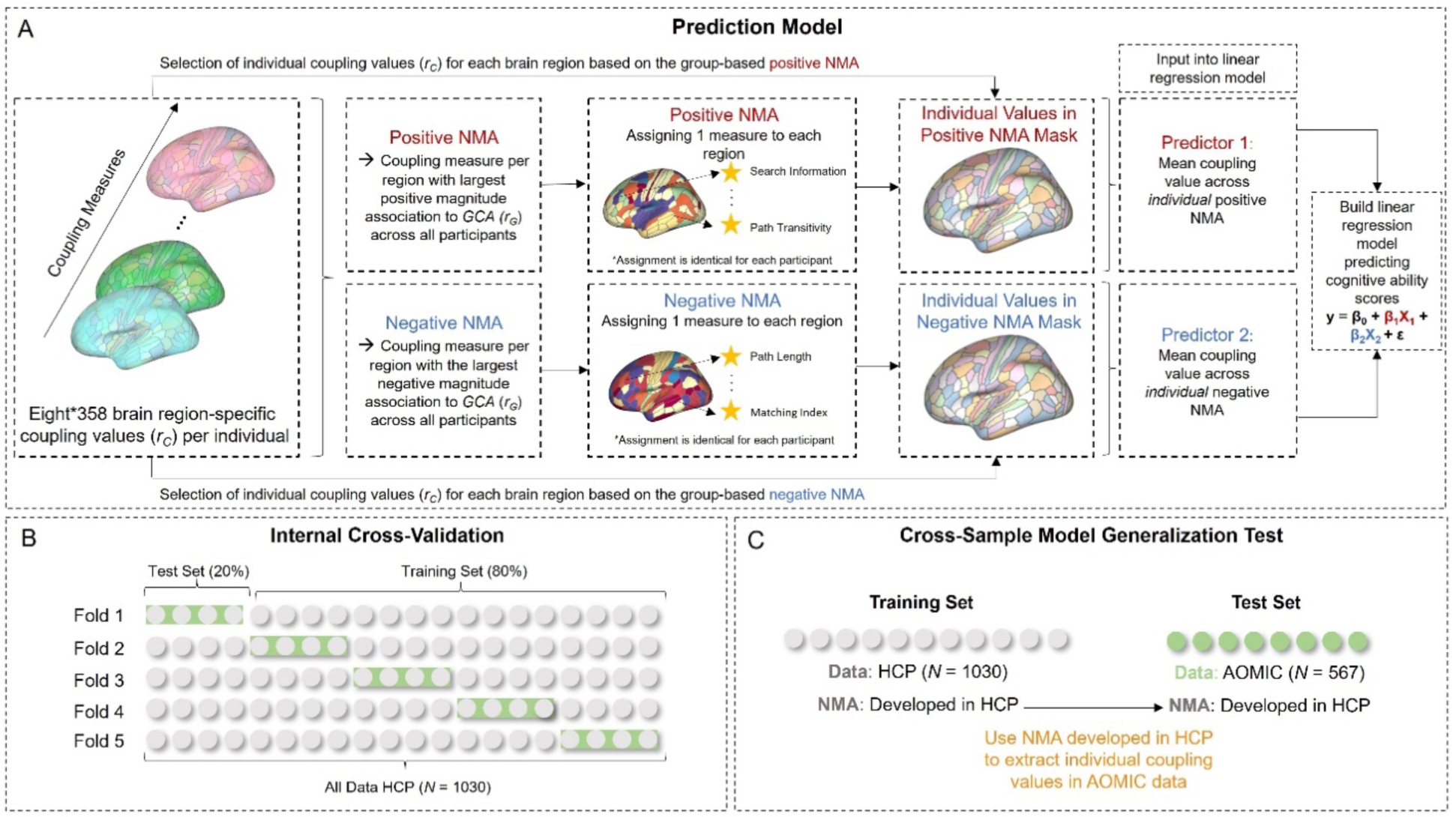
Workflow of the predictive modeling framework used to predict individual cognitive ability scores from region-specific SC-FC coupling. (A) In general, prediction models were built using two input predictor variables that were derived from individual’s coupling values (r_C_) extracted by using group-based positive and a negative node-measure assignment (NMA) masks. For the creation of the group-based positive and negative NMA masks, the coupling measure with the largest positive and negative magnitude associations between coupling measures with GCA (r_G_) per brain region across all participants was selected. These group-based NMA masks defined which individual-specific coupling values were chosen for each brain region and the two predictors for the linear regression model were computed by taking a brain-average across each individual’s positive NMA (predictor 1) and negative NMA (predictor 2). (B) For the 5-fold internal cross-validation, the model was trained on four folds (80%; *N_train_* ∼ 825) of the sample and then used to predict cognitive ability scores in the withheld fold (20%; *N_test_* = ∼ 205) of the sample. To avoid data leakage between cross-validation folds, the group-based positive and negative NMA masks built in the training sample were used to extract individual coupling values in the respective test sample. This procedure was repeated five times and prediction accuracy was assessed by correlating predicted and observed cognitive ability scores. Equal distribution of subjects in folds with respect to GCA and family relations was guaranteed by using stratified folds. (C) For the cross-sample model generalization test, the prediction model was built using data from the complete main sample (HCP) and then tested on an independent sample (replication sample, AOMIC). Here, group-based positive and negative NMA masks were generated based on data from the complete main sample. HCP = Human Connectome Project; AOMIC = Amsterdam Open MRI Collection; NMA = Node-Measure Assignment; GCA = General Cognitive Ability. [color; 2-column fitting image]

To assess for visualization purposes the coupling strength corresponding to the positive and negative NMA masks, brain region-specific coupling values in the positive and negative individual NMAs were separately averaged across all participants, thus yielding a group-average map of regional coupling values (i.e., coupling strength) for the positive and negative NMA, respectively.

#### 2.11.2. Cross-validated prediction framework

A 5-fold cross-validated multiple linear regression model was implemented to predict individual cognitive ability scores. Specifically, the sample was split into five folds by simultaneously ensuring equal distributions of subjects with respect to GCA (via stratified folds) and by accounting for family relations. Then, the model was trained on 80% of the sample (*N_train_* ∼ 825) and afterwards used to predict cognitive ability scores in the withheld 20% of the sample (*N_test_* = ∼ 205). This procedure was repeated five times so that every subject was part of the test set once and received one predicted cognitive ability score (Fig. 2B). The predictor variables of the linear regression model:

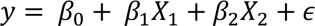

were a) brain-average coupling values from the individual positive NMAs (*X_1_*) and b) brain-average coupling values from the individual negative NMAs (*X_2_*), where *β*_0_ is the y-intercept, *β_n_* is the slope (or regression coefficient), *ε* is the error term and *y* the predicted intelligence score. Note that we used the positive and negative NMAs built in the training sample to extract individual coupling values in the respective test sample. This step is important to keep the training and test sets strictly independent from one another and to avoid leakage of information between folds. All input parameters were normalized before model building, and confounding variables (age, gender, handedness, and in-scanner head motion) were regressed out with linear regression from all variables (brain-average coupling values from the individual positive and negative NMAs; cognitive ability scores) for each training sample. The regression model of the training sample was then applied to the test sample of the respective fold.

As a result of the 5-fold cross-validated prediction, each individual was part of the test set once and thus received a predicted cognitive ability score. Model performance was assessed by correlating observed and predicted cognitive ability scores across the whole sample. The 5-fold cross-validated prediction was repeated 100 times with different training-test data splits and prediction performance was averaged across all 100 runs. To assess the significance of the prediction, a non-parametric permutation test with a total of 1000 iterations as recommended by Dubois et al. (2018) was performed. Specifically, individual cognitive ability scores were permuted, and models were trained with these permuted scores by applying exactly the same approach as outlined above. Performance of the model trained on the permuted cognitive ability scores was then compared to model performance using the true values. This procedure was conducted 10 times for each different training-test data partition, yielding 1000 total repetitions. The *p*-value indicating statistical significance was calculated by evaluating how often the model trained on the permuted scores was better at predicting the observed scores.

### 2.12. External replication

To test the robustness of our results, all analyses were repeated in a completely independent sample that differs in the measure of GCA, data acquisition and preprocessing (AOMIC, Snoek et al., 2021). Specifically, we used data from the ID1000 sample (*N* = 928) consisting of healthy subjects of age 19-26 (483 female, 826 right-handed, mean age = 22.8 years). All MRI data were acquired with a gradient-echo EPI sequence on a Philips Intera 3T scanner with a 32-channel head coil. Three diffusion-weighted scans and one functional (BOLD) MRI scan were recorded. Importantly, in this sample no resting-state fMRI scan was conducted as participants were passively watching a movie clip consisting of uneventful natural scenes that has been demonstrated to allow for good approximation of intrinsic connectivity (Vanderwal et al., 2017). FMRI scans were obtained without multi-slice acceleration (TR = 2200 ms, TE = 28 ms, 3-mm isotropic voxel resolution, and flip angle = 90°). The fMRI data were downloaded in the minimally preprocessed form and further preprocessed similarly as in the main sample. Structural imaging data were acquired from diffusion weighted imaging (median TR = 6312 ms, TE = 74 ms, 2-mm isotropic voxel resolution, b = 1000 s/mm^2^, and 32 directions/shell) and preprocessed using the same steps applied to the HCP data to model white matter fibers but using a maximum spherical harmonics order of six. Further details on image acquisition and preprocessing are described in Snoek et al. (2021). GCA was operationalized with an established intelligence measure, i.e., the Intelligence Structure Test (IST), assessing verbal, numerical and figural abilities (Beauducel et al., 2010). More specifically, the summed scores of the three measures crystallized intelligence, fluid intelligence and memory were used. After subject exclusion based on missing demographic, neuroimaging or behavioral data, low quality of structural images and motion exclusion based on framewise displacement (same criteria as in the main sample), 567 subjects remained in the replication sample (age 19-26, 300 female, 500 right-handed, mean age = 22.8 years). The following analyses were repeated in the replication sample: a) SC-FC coupling operationalized with similarity measures and communication measures, b) investigation of associations between brain-average SC-FC coupling and cognitive ability scores, c) investigation of associations between region-specific SC-FC coupling and cognitive ability scores with an internally cross-validated prediction framework. Individual raw IST-scores were used for the correlative analyses between GCA and measure-specific brain-average SC-FC coupling values. For the predictive analyses, IST scores were normalized.

Finally, a cross-sample model generalization test was conducted (Fig. 2C). Specifically, we evaluated whether our prediction model that accounts for region-specific variations in the type of SC-FC coupling built in the main sample (HCP) could also predict individual cognitive ability scores in the replication sample (AOMIC). Model performance was evaluated by correlating the predicted with the observed cognitive ability scores. Significance of the prediction was assessed with a permutation test. Specifically, cognitive ability scores in the main sample (HCP) were permuted and the model was trained with these permuted scores. Prediction performance of the model trained on the permuted scores was then compared to model performance using the true values. This procedure was repeated 1000 times. The *p*-value indicating statistical significance was calculated by evaluating how often the model trained on the permuted values was better at predicting the observed scores.

### 2.13. Ethical approvals

For the main sample, all study procedures were approved by the Washington University Institutional Review Board (details see Van Essen et al., 2013). For the replication sample, study procedures were authorized by the Ethics Review Board of the department of Psychology at the University of Amsterdam (details see Snoek et al., 2021). Written informed consent in accordance with the declaration of Helsinki was obtained from all participants of the main sample and the replication sample.

### 2.14. Data Availability Statement

All analyses were implemented in Python (version 3.4) and MATLAB (Version R2021a). Data of the main sample can be accessed under https://www.humanconnectome.org/study/hcp-young-adult/data-releases/ and data from the replication sample can be obtained under https://openneuro.org/datasets/ds003097. All analysis code for the current study is available on GitHub: DWI preprocessing: https://github.com/civier/HCP-dMRI-connectome; fMRI preprocessing: https://github.com/faskowit/app-fmri-2-mat; computation of latent *g*-factor: https://github.com/jonasAthiele/BrainReconfiguration_Intelligence; operationalization of SC-FC coupling with communication measures: https://github.com/brain-networks/local_scfc; main analysis and replication analysis as implemented in the current study: https://github.com/johannaleapopp/SC_FC_Coupling_Cognitive_Ability.

## 3. Results

### 3.1. General cognitive ability and SC-FC coupling

Individual GCA was operationalized as latent *g*-factor from 12 cognitive scores with bi-factor analysis (Dubois et al., 2018; Thiele et al., 2022) using data from 1086 subjects from the HCP (Van Essen et al., 2013). In the replication sample (AOMIC), cognitive ability was assessed with the Intelligence Structure Test (IST, Beauducel et al., 2010). Both measures were approximately normally distributed (see Supplementary Fig. S1). In the HCP, the *g*-factor ranged between −2.60 and 2.40 (*M* = 0.08; *SD* = 0.89) and in the replication sample, the sum score of the IST ranged between 78 and 295 (*M* = 202.85; *SD* = 39.01).

Individual-specific SC-FC brain network coupling was operationalized with two similarity measures and six communication measures (computed based on each individual SC matrix) that were set in relation to each individual’s FC, resulting in 358 individual- and region-specific coupling values (*r_C_*) for each of the eight coupling measures (see Methods and Fig. 1). A group-average SC-FC coupling map (Fig. 3) was calculated by first identifying the similarity or communication measure per brain region that was able to most frequently explain the highest amount of variance (*R^2^*) in the respective region’s functional connectivity profile across all participants. Subsequently, this group-general mask was used to extract individual brain region-specific coupling values (*R^2^*) that were then averaged across subjects (see Methods). In line with previous reports (Baum et al., 2020; Griffa et al., 2022; Gu et al., 2021; Vázquez-Rodríguez et al., 2019; Zamani Esfahlani et al., 2022), highest coupling was observed in somatomotor and visual areas (average maximal *R^2^* ∼ .3), while lowest coupling was identified in parietal and temporal regions (average maximal *R^2^* ∼ .075). Note that the here used approach (selecting measures based on highest coupling strength) allowing for individual variability in the measure chosen per brain region was solely used for visualization purposes (Fig. 3, Supplementary Fig. 3, and Supplementary Fig. 8), but not for further statistical analyses. In the subsequent correlative and predictive analyses, region-specific coupling measures were selected based on the strongest association with GCA by using cross-validated group-general masks (the positive NMA and the negative NMA).

**Fig. 3.**
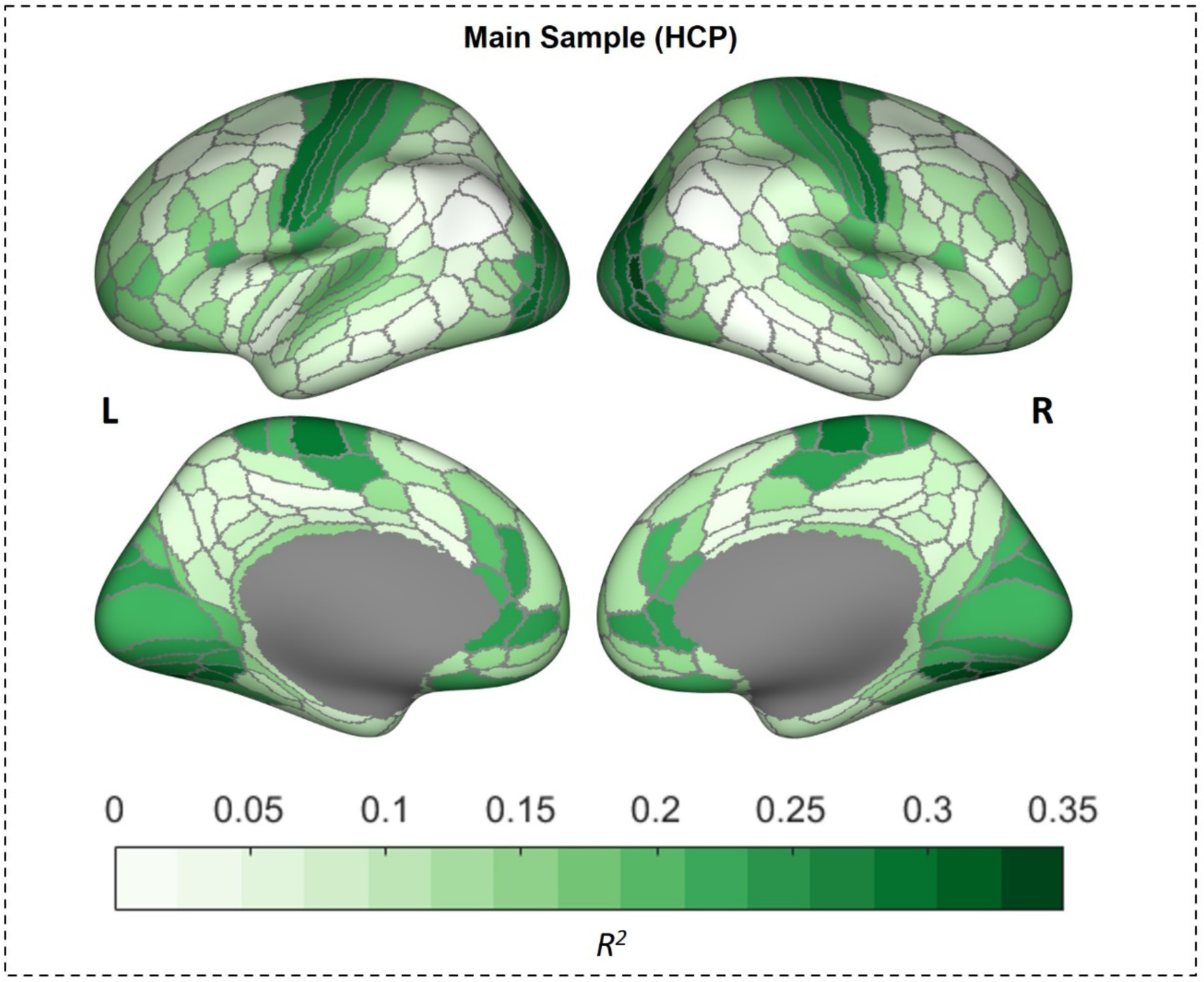
Group-average whole-brain pattern of SC-FC coupling strength. This figure illustrates the cortical distribution of coupling strength based on the region-specific similarity or communication measure (computed based on structural connectivity) able to explain the highest amount of variance in functional connectivity most frequently across all participants in the main sample (HCP). This group-general mask was subsequently used to extract individual region-specific coupling values (*R^2^*) and a group-average map of the regional coupling strength was created by averaging across all participants’ coupling values. Note that this approach was solely used for visualization purposes but not for further statistical analyses. In correlative and predictive analyses, the coupling measure was determined by a group-general mask (i.e., the positive and negative NMAs) based on the strongest association with GCA. These latter masks were also cross-validated in all predictive approaches to prevent any data leakage between training and test samples. [color; single column fitting image]

### 3.2. General cognitive ability is associated with brain-average SC-FC coupling operationalized with path transitivity

GCA was significantly associated with brain-average SC-FC coupling (subject-specific average across all regional coupling values per coupling measure) operationalized with the communication measure path transitivity (*r* = .10, *p* = .002, corrected for multiple comparisons). All other associations between brain-average coupling and GCA were of negligible effect sizes and did not reach statistical significance (all *r* < .09, all *p* > .006; Table 3).

**Table 3.** Relationship between general cognitive ability and brain-average SC-FC coupling.

| Measure to Compute SC-FC Coupling | Main Sample – HCP $r(p)$ |
| --- | --- |
| Cosine Similarity (CoS) | .05 (.115) |
| Matching Index (MI) | .07 (.031) |
| Communicability (G) | .04 (.247) |
| Mean First Passage Time (MFPT) | .03 (.415) |
| Flow Graphs (FG) | .08 (.010) |
| Search Information (SI) | .02 (.588) |
| Path Transitivity (PT) | .10 (.002)* |
| Path Length (PL) | .05 (.095) |

*Note:* Main sample $N = 1030$ . Partial correlations between cognitive ability scores and individual measure-specific brain-average coupling (measure-specific average of coupling values from all brain regions) controlled for age, gender, handedness, and in-scanner head motion. Significant associations passing the Bonferroni-corrected threshold (eight comparisons) are marked with an asterisk (\* = $p < .006$ ).

### 3.3. The relation between general cognitive ability and SC-FC coupling varies between different brain regions

As previous research suggests that preferred communication strategies differ between brain regions (Betzel et al., 2022; Zamani Esfahlani et al., 2022), we next developed a cross-validated prediction framework that a) allows to examine the relationship between GCA and region-specific SC-FC coupling, b) avoids the multiple comparisons problem, and c) tests if individual cognitive ability scores can be predicted by region-specific SC-FC coupling. Features for the prediction model were individuals’ brain region-specific coupling values (*r_C_*) that were selected by applying group-average node-measure assignment (NMA) masks. For the internally cross-validated prediction model, we computed for each training sample one NMA mask with the largest positive and one NMA mask with the largest negative magnitude associations between coupling measures and GCA (*r_G_*; see Methods and Fig. 2A). These training sample-based masks were used for all subjects of the corresponding test sample to extract individual-specific coupling values that were then averaged separately for the positive and negative individual NMA and represent the model features. For the cross-sample model generalization test, additional NMA masks were generated based on data from the complete main sample (*N* = 1030). These latter masks are illustrated in Fig. 4A and confirm that the GCA-associated type of coupling (as indicated by different coupling measures) as well as the associated coupling strength (Fig. 5) varies across the cortex. For the positive NMA, highest coupling was observed in frontal areas, while lowest coupling was identified in somatomotor and visual areas. For the negative NMA, highest coupling was detected in somatomotor and visual areas, while lowest coupling was observed in parietotemporal areas. Notably, as region-specific measures (ultimately defining the prediction model features, i.e., the individual coupling values that were extracted based on these measures) were selected based on their association with general cognitive ability, the creation of NMA masks served as means to investigate which coupling measure was most or least able to account for individual differences in GCA.

**Fig. 4.**
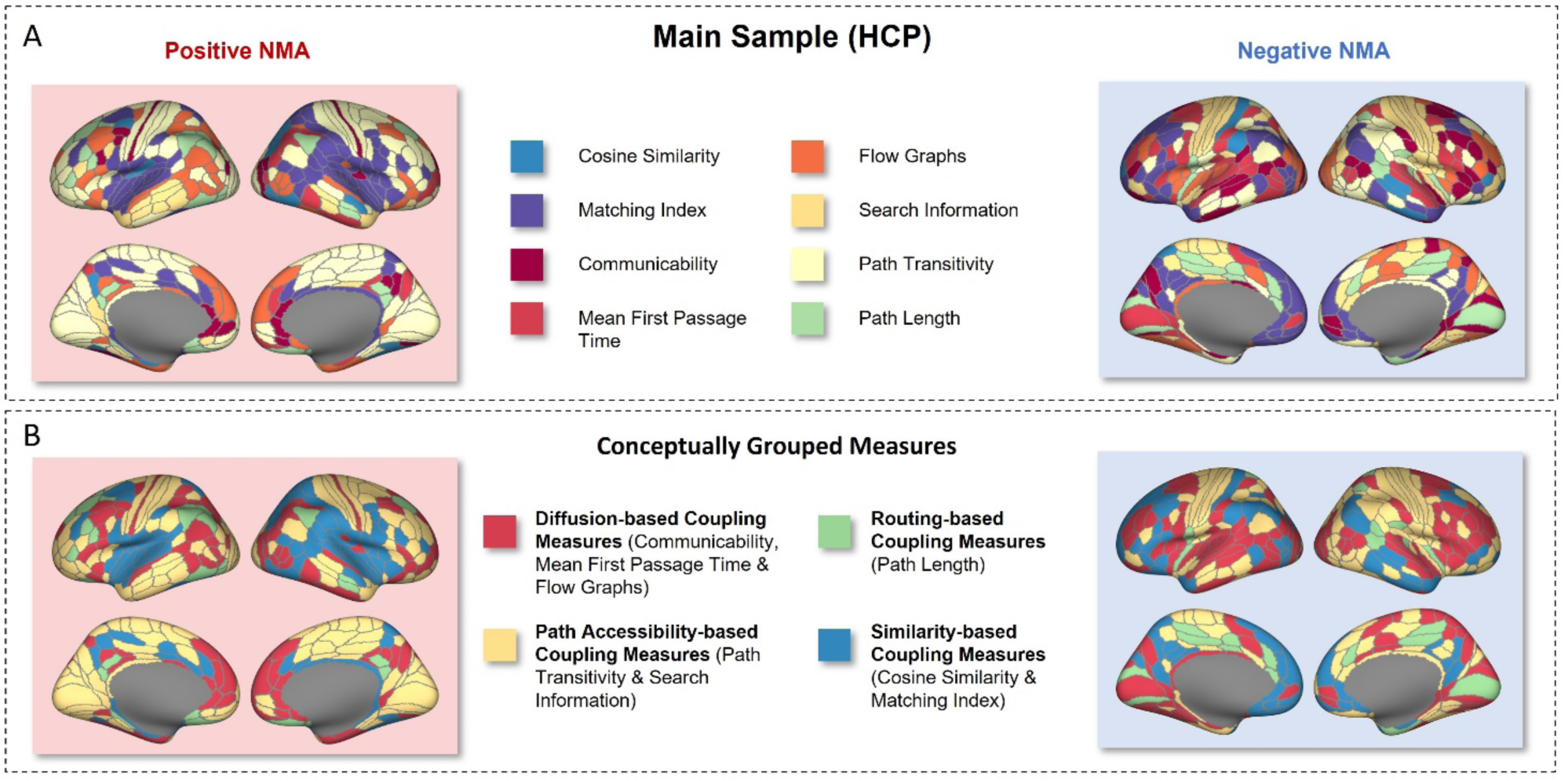
General cognitive ability is associated with brain region-specific SC-FC coupling. (A) Group-based positive and negative node-measure assignment (NMA) masks as used in the cross-sample model generalization test. These were created by identifying the coupling measure with the largest positive and negative magnitude associations with GCA (*r_G_*) per brain region across all participants of the main sample (*N* = 1030), depicting the measure per brain region best explaining individual differences in GCA. Note that to thoroughly prevent overfitting, for the 5-fold internal cross-validation, NMAs were created separately for each training sample of each cross-validation loop and could thus slightly differ from the whole sample-based masks shown in this illustration. Individual-specific coupling values (*r_C_*) were then extracted with these masks, i.e., the group-based NMA mask defined which individual-specific coupling value (one out of eight different coupling values, see Methods) was extracted for each brain region and used for further analyses. (B) For illustration purposes, coupling measures were grouped based on conceptual similarity of the proposed signaling mechanism. The grouped positive NMA mask revealed that coupling measures based on similarity were chosen predominantly in dorsotemporal regions and coupling measures based on path accessibility were selected primarily for frontal regions, but also for a widely distributed set of brain regions. The grouped negative NMA mask showed that coupling measures based on similarity were only very seldomly selected, while coupling measures based on diffusion were chosen for many brain regions and especially frequently in frontal brain regions. HCP = Human Connectome Project; NMA = Node-Measure Assignment. [color; 1.5-column fitting image]

**Fig. 5.**
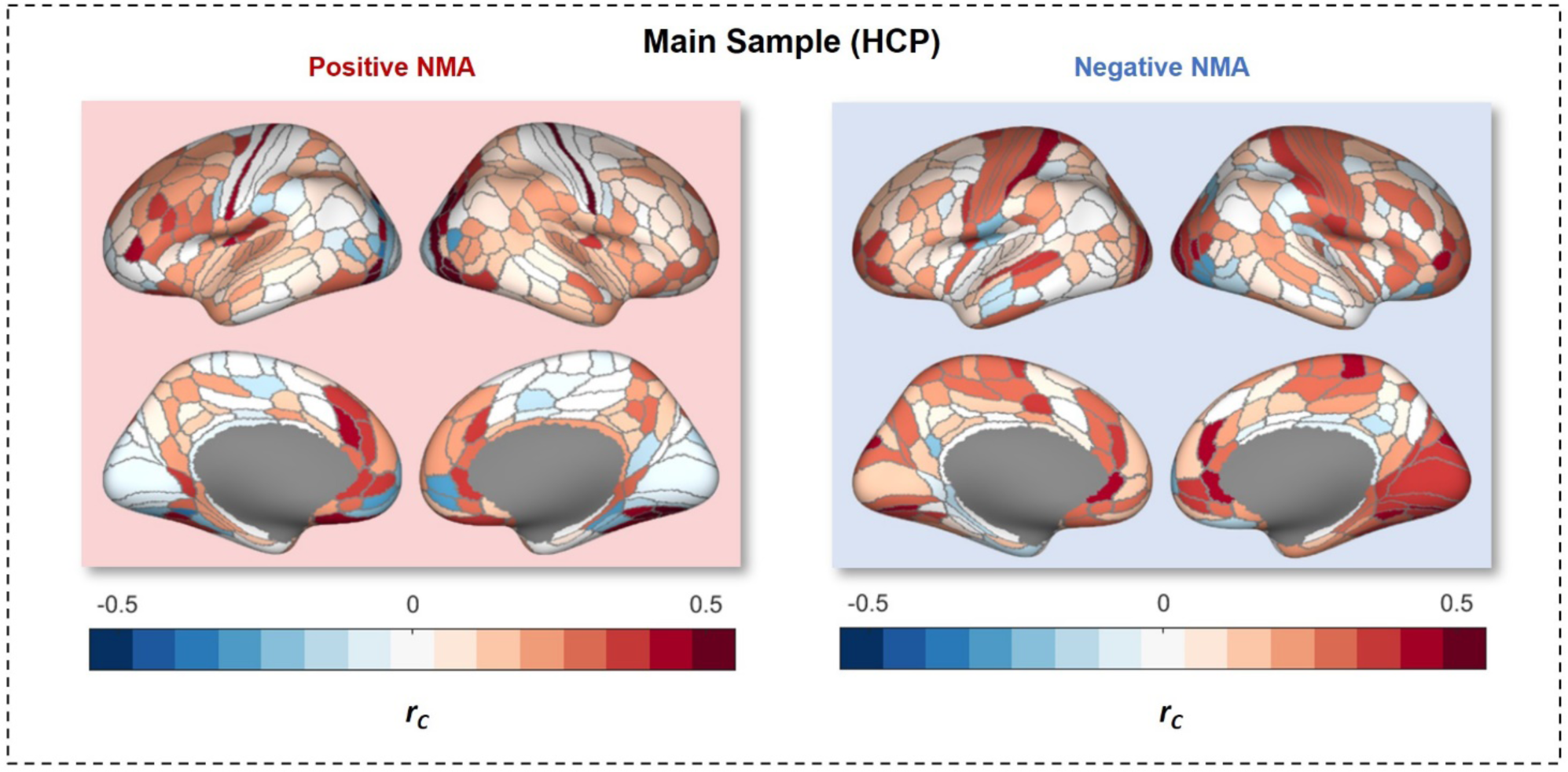
Group-average region-specific SC-FC brain network coupling strength corresponding to the node-measure assignment (NMA) masks as used in the cross-sample model generalization test. These were created by identifying the coupling measure with the largest positive and negative magnitude associations with GCA (*r_G_*) per brain region across all participants of the complete main sample (*N* = 1030). For each participant, individual-specific coupling values were extracted using the group-based NMA masks, resulting in two individual brain maps (for the positive and negative NMA mask, respectively) containing one coupling value (*r_C_*) for each brain region. This figure illustrates a group-average map of these individual regional coupling values. HCP = Human Connectome Project; NMA = Node-Measure Assignment. [color; 1.5-column fitting image]

To better visualize the distribution of coupling measures in the NMA masks, the eight coupling measures were grouped based on conceptual similarity into a) diffusion-based coupling measures (communicability, mean first passage time, flow graphs), b) coupling measures based on path accessibility (search information, path transitivity), c) routing-based coupling measures (path length), and d) coupling measures based on similarity (cosine similarity, matching index). The grouped positive NMA mask revealed that coupling measures based on similarity of the structural connectivity profiles were selected predominantly in dorsotemporal regions and coupling measures based on path accessibility were selected primarily for frontal regions, but also for a widely distributed set of brain regions (Fig. 4B). The grouped negative NMA revealed that coupling measures based on similarity were only very seldomly chosen, while coupling measures based on diffusion were selected for many brain regions and especially frequently in frontal brain regions (Fig. 4B). The spatial relation of GCA-associated coupling measures for the positive and negative NMA to the seven functional network partition of Yeo et al. (2011) is listed in Supplementary Table S1.

### 3.4. Region-specific SC-FC coupling predicts general cognitive ability scores in unseen individuals

The prediction model based on linear regression with two predictors, i.e., brain-average across each individual positive NMA (predictor 1) and across each individual negative NMA (predictor 2; see Methods), succeeded to significantly predict individual cognitive ability scores (5-fold cross-validation; correlation between observed vs. predicted cognitive ability scores: *r* = .25, *R^2^* = .06*, p <* .001 by permutation test). A scatterplot depicting the relationship between predicted and observed cognitive ability scores is illustrated in Supplementary Fig. S2A. For completeness, the performances of this model to predict individual cognitive performance scores (see Table 1) that were used to compute the latent *g*-factor are listed in Supplementary Table S2.

### 3.5. External replications suggest robustness of findings and model generalizability

To ensure that our research findings are robust and generalizable to the population, all analyses were replicated in an independent sample, i.e., the AOMIC ID1000 cohort (*N* = 567, Snoek et al., 2021). Similar as in the main sample, general cognitive ability was significantly positively associated with brain-average coupling operationalized with path transitivity (*r* = .16, *p* < .001). Again, all other brain-average coupling measures were not significantly correlated with GCA (Supplementary Table S3). The group-average whole-brain pattern of SC-FC coupling strength is illustrated in Supplementary Fig. S3 and brain maps visualizing the positive and negative NMAs are shown in Supplementary Fig. S4. Coupling strengths corresponding to the NMAs are visualized in Supplementary Fig. S5. The 5-fold cross-validated multiple linear regression model with input features created with the same strategy as in the main sample (see Methods) was also able to significantly predict individual cognitive ability scores within the replication sample (correlation between observed vs. predicted scores *r* = .17*, R^2^* = .03, *p* = .002 by permutation test, Supplementary Fig. S6).

Model generalizability was tested with a cross-sample model generalization test. To this aim, the complete main sample (*N* = 1030) was used for model building and the prediction model was then tested with input data from the replication sample (*N* = 567, Fig. 2C). The model based on the HCP data was able to significantly predict individual cognitive ability scores also in the AOMIC sample (correlation between observed vs. predicted scores *r* = .19*, R^2^*= .04, *p* < .001 by permutation test; Supplementary Fig. S2B).

### 3.6. Post-hoc analyses

Several post-hoc analyses were performed (on the main sample only) to further evaluate the robustness of our findings and to enhance their interpretation.

At first, since it is an established finding that GCA is significantly positively associated with total brain volume (see e.g., McDaniel, 2005; Pietschnig et al., 2015), and individual cognitive ability scores indeed correlated significantly with total intracranial volume also in our sample (*r* = .33, *p* < .001), all analyses were repeated with total intracranial volume as additional control variable. Correlative results investigating the relationship between brain-average coupling and GCA were highly similar (Supplementary Table S4). Also, the correlation between predicted and observed cognitive ability scores in the 5-fold cross-validated prediction framework remained significant (*r* = .21, *R^2^* = .04, *p* < .001 by permutation test).

Second, the group-average SC-FC coupling pattern (Fig. 3) indicates overall higher SC-FC coupling in brain areas associated with unimodal neural processing (somatomotor and visual areas) and lower coupling in areas associated with multimodal processing (frontal, parietal and temporal areas). Further, the region-specific assignment of coupling measures in the group-average node-measure assignment masks (NMAs; Fig. 4) also points to differences between unimodal and multimodal brain areas. To evaluate this pattern statistically, a principal gradient referred to as Margulies gradient (specified by Margulies et al., 2016; Supplementary Fig. S7), which situates each brain region on a spectrum between unimodal regions serving primary sensory and motor tasks (negative values) and multimodal regions serving complex heteromodal activity (positive values), was tested for its numerical association with a) the group-average region-specific SC-FC coupling strength (Fig. 3) and b) the assignment of coupling measures in the positive and negative NMA mask (Fig. 4). The vector describing the group-average region-specific SC-FC coupling strength (*R^2^*) and the Margulies gradient were significantly negatively correlated (Pearson correlation; *r* = −.45, *p* < .001 by spin permutation testing), thus supporting the post-hoc assumption that higher SC-FC coupling is more likely to be present in unimodal brain areas while lower SC-FC coupling is predominant in multimodal brain areas. Associations with the pattern of coupling measure assignment in the positive and negative NMA were analyzed by comparing Margulies gradient values from brain regions of the four coupling measure groups (diffusion, path accessibility, routing and similarity; Fig. 4B) with a one-way ANOVA and Tukey’s HSD test for multiple comparisons (Supplementary Table S5). Results indicate that in the positive NMA, routing-based measures were predominantly selected in multimodal areas (i.e., positive group-average of Margulies gradient values) while similarity-based measures were frequently chosen in unimodal areas (i.e., negative group-average of Margulies gradient values). In the negative NMA, similarity-based measures were selected in multimodal areas while routing-based measures were chosen in unimodal areas.

Third, it was ensured that the regression of global signals applied in the fMRI preprocessing did not affect the results. Therefore, a control analysis was implemented where SC-FC coupling was operationalized similarly as in the main analysis, but fMRI preprocessing was conducted without global signal regression (see Parkes et al., 2018, strategy no. 7). Results were very similar between both preprocessing strategies (correlation between the two vectors depicting the brain region-specific SC-FC coupling strength with and without global signal regression: *r* = .98, *p* < .001; see Supplementary Fig. S8). Group-based positive and negative NMA masks were also re-computed across the complete sample. Supplementary Fig. S9 highlights that the NMAs were nearly identical. Finally, the correlation between predicted and observed cognitive ability scores in the 5-fold cross-validated prediction model developed based on information from region-specific SC-FC coupling with or without global signal regression was comparable: *r* = .25 vs. *r* = .23, both *p* < .001 by permutation test. Overall, these results suggest that applying global signal regression during fMRI preprocessing does neither affect the region-specific SC-FC coupling pattern significantly nor does it influence the associations with general cognitive ability markedly.

Fourth, we tested which particular features drove the performance of the prediction model. To answer the question if there exists a single communication measure underlying significant prediction, we reran the internally cross-validated prediction model in the main sample and excluded one measure from the selection process (creation of positive or negative NMA) at a time. Prediction performances (predicted vs. observed general cognitive ability scores) were all in a similar range (*r* = .21 - .26; Supplementary Table S6) revealing that there exists no particular coupling strategy that outweighs the others. Additionally, we were interested in the question whether the significant performance of the prediction model depends on the positive network feature, the negative network feature, or on both. Therefore, we reran the internally cross-validated prediction model in the main sample while excluding one of the two features at a time (Supplementary Table S7). Only using information from coupling measures that are positively associated with GCA (positive network feature only) still enabled significant prediction of individual general cognitive ability scores (correlation between observed vs. predicted scores *r* = .21; *p* = .001), while only using information from coupling measures negatively associated with general cognitive ability (negative network feature only) was not sufficient to significantly predict individual differences in GCA (*r* = .03; *p* = .238). However, combining both features generated the best prediction performance (*r* = .25; *p* < .001) and reveals most about the relationship of SC-FC coupling and GCA expanding on previous reports (see e.g., Baum et al., 2020).

As outliers in the region-specific SC-FC coupling data could have potentially influenced the performance of the prediction model, especially because features were constructed by taking an average across all values, we lastly implemented two additional control analyses to ascertain that this was not the case. To get a first impression of the quantity of outliers (defined as elements > 3SD away from the mean) in region-specific coupling values, the NMA masks computed across the complete sample (see Fig. 4) were used to extract individual values. Across all individual region-specific coupling values (1030*358) in the positive NMAs, the percentage of outliers with respect to other coupling values from the same brain region was 0.42% while in the negative NMAs, the respective percentage was 0.36%. These percentages are in line with what would be expected in normally distributed data. Nevertheless, we tested for any potentially remaining impacts of outliers on prediction model performance by rerunning the internally cross-validated prediction model whilst eliminating outliers in training sample-specific and test sample-specific individual NMAs before computing the individual average values that were used as model features. As expected, removing these outliers did not have a noticeable influence on prediction performance (correlation between observed vs. predicted scores *r* = .25; *p* < .001).

In sum, the post-hoc control analyses suggest that a) our findings were not confounded by individual differences in total brain volume, b) the overall SC-FC coupling pattern and the assignment of coupling measures in the NMAs both correspond to the unimodal-multimodal macroscale cortical organization as defined by the Margulies gradient (Margulies et al., 2016), c) the here reported results do not depend on whether or not global signal regression was applied during the preprocessing of the fMRI data, d) the significant prediction of GCA scores from region-specific SC-FC coupling information does not depend on a single coupling measure, but the positive network feature is mostly driving the significant prediction, and e) that the performance of the prediction model is not noticeably influenced by outliers in the region-specific SC-FC coupling data.

## 4. Discussion

The aim of this study was to investigate if individual variations in structural-functional brain network coupling (SC-FC coupling) are associated with individual differences in general cognitive ability (GCA). We used data from the Human Connectome Project (HCP; *N* = 1030) and operationalized SC-FC coupling with two similarity measures and six communication measures. At the whole-brain level, higher GCA was associated with stronger SC-FC coupling, but only for path transitivity as communication strategy. By focusing on brain region-specific variations in coupling measures and by accounting for positive and negative associations with GCA, we showed that individual cognitive ability scores can be predicted from SC-FC coupling within a cross-validated prediction framework. Notably, all analyses were replicated in an independent sample and the prediction model built in the main sample also succeeded to significantly predict cognitive ability scores in the replication sample (*N* = 567), together suggesting high robustness of study findings and cross-sample generalizability of the prediction model.

### 4.1. SC-FC coupling has a unique distribution across the human cortex

The spatial pattern of group-average SC-FC coupling observed in our study highly resembled the distribution reported in prior studies, i.e., highest coupling was observed in visual and somatomotor areas and lowest coupling in parietal and temporal areas (Baum et al., 2020; Gu et al., 2021; Vázquez-Rodríguez et al., 2019; Zamani Esfahlani et al., 2022). In addition to previous reports, our post-hoc analysis revealed that higher coupling was prevalent in unimodal areas while lower coupling was predominant in multimodal areas. Differences in microstructure (e.g., intracortical myelination and laminar differentiation) along the gradient spanning from unimodal to multimodal areas (Huntenburg et al., 2017; Margulies et al., 2016; Paquola et al., 2019; Vázquez-Rodríguez et al., 2019), which are thought to be rooted in the rapid evolutionary expansion of the human cortex (Buckner and Krienen, 2013), might represent one possible explanation for this observation. More specifically, it can be speculated that the untethering of brain structure and function in multimodal areas, as indexed by lower SC-FC coupling, results from the frequent reconfiguration of local microcircuitry and overall less signaling constraints as required for polysensory integration (Buckner and Krienen, 2013; Vázquez-Rodríguez et al., 2019).

### 4.2. SC-FC coupling via path transitivity is associated with general cognitive ability

At the whole-brain level, variations in SC-FC coupling operationalized with the communication measure path transitivity were positively associated with individual differences in GCA. On the one hand, the extent to which two brain regions show synchronized activity (i.e., are functionally connected) is thought to be related to the ease of which neural signals can propagate based on the underlying structural connections (Goñi et al., 2014). On the other hand, path transitivity measures this ease of communication by reflecting the accessibility of the shortest structural path based on the number of available detours carrying neural signals back to the shortest path when the direction is lost. Higher correspondence between path transitivity derived from structural connectivity and functional connectivity could therefore imply that signal transmission, leading to functional interactions between two brain regions, is operating more closely along transitive structural paths. The availability of local detours, quantified by path transitivity, may counteract signal dispersion and create feedback loops to recurrently stabilize signals and thus enable signals to re-access the shortest path after having left, which may support efficient communication (Goñi et al., 2014). From a more theoretical point of view, one of the most popular neurocognitive theories of intelligence differences, the Neural Efficiency Hypothesis of Intelligence (NEH, Haier et al., 1988; Neubauer and Fink, 2009) explicitly assumes that people with higher intelligence scores require less brain activation while performing cognitive tasks and are therefore more capable of efficient neural processing (Dunst et al., 2014; Neubauer and Fink, 2009). The results reported here inform this theory by proposing how efficient neural processing in individuals with higher cognitive ability may depend on interactions between brain structure and brain function.

### 4.3. Regional specificity in communication strategies facilitates cognition

The finding that communication measures associated with GCA vary critically between different brain regions complements previous research proposing that information integration in the brain is not just facilitated by one unique signaling mechanism but that regional variability of communication strategies is evident (Betzel et al., 2022), that it improves the prediction of individual functional connectivity (Zamani Esfahlani et al., 2022) and ultimately of human traits (Seguin et al., 2020). The regional variability of preferred communication strategies can also be interpreted against the background of psychological theories and recent studies indicating that individual variation in the coordinated action of several cognitive processes including working memory, long-term memory, cognitive flexibility, and processing speed underlies differences in general cognitive ability (Duncan et al., 2020; Frischkorn et al., 2019; Guilford, 1967; Kovacs and Conway, 2016; Neisser et al., 1996). More specifically, visual inspection of the node-measure assignments and the results of post-hoc analyses suggest that in multimodal areas that have frequently been associated with such higher-order cognitive functions (Betzel et al., 2022; Dosenbach et al., 2007), a higher prevalence of directed communication strategies (coupling measures based on routing and path transitivity) is associated with higher GCA. In contrast, in unimodal areas a lower prevalence of directed communication strategies was associated with higher GCA (Supplementary Table S5). More directed signaling strategies operating along the shortest path could contribute to faster and more efficient information integration in multimodal areas, while the disadvantage of directed signaling strategies in unimodal areas needs to be clarified by future research, e.g., whether directed processes might not always be possible and thus diffusive processes would be preferable, especially when final destinations of the signals are unknown. Nevertheless, our results support that both such effects, i.e., higher prevalence of directed communication strategies in multimodal areas and lower prevalence of directed communication strategies in unimodal brain areas facilitate higher GCA.

### 4.4. Cross-validation and independent replication support robustness and generalizability of findings

Successful prediction of individual cognitive ability scores based on the combination of information from multiple brain areas reinforces established neurocognitive models of intelligence including the Parieto-Frontal Integration Theory (P-FIT, Jung and Haier, 2007; Basten et al., 2015) and the Multiple Demand Theory (Duncan, 2010) as well as recent investigations (Barbey, 2018; Hilger et al., 2020; Thiele et al., 2022) suggesting that a distributed network of brain regions underlies individual differences in GCA. From a methodological point of view and especially important against the background of the replication crisis in psychological science (Open Science Collaboration, 2015; Poldrack et al., 2020, 2017), predictive approaches using two independent samples for model building and model testing reduce the danger of overestimating effect sizes by fitting sample-specific variance, as often the case in correlative approaches (overfitting; Cwiek et al., 2022; Yarkoni and Westfall, 2017). Latest guidelines critically assessing machine learning practices in neuroimaging emphasize the particular importance of replication in an external and thus completely independent dataset compared to internal cross-validation (Cwiek et al., 2022; Marek et al., 2022). Repeated subsampling within the same dataset (internal cross-validation) often still overestimates the reproducibility of effects, due to variance that is shared between sub-samples. This shared variance is, however, not attributable to the same scaling on the variable of interest (e.g., general cognitive ability) but rather originates from the application of the same methodology during e.g., preprocessing or the existence of common patterns of biases inherent to the whole dataset (Poldrack et al., 2020; Tervo-Clemmens et al., 2023). To circumvent this pitfall and to best demonstrate generalizability of study results, independent out-of-sample evaluation seems to be the best solution (Poldrack et al., 2020; Tervo-Clemmens et al., 2023). Our study that combines internal cross-validation, the replication of findings in an independent sample as well as a cross-sample model generalization test, follows these best practices. Being able to successfully predict individual GCA scores in the independent replication sample, despite different imaging states and general cognitive ability measures, implies high robustness of findings as well as generalizability of the here developed prediction model.

## 5. Limitations and future directions

Several limitations need to be mentioned. At first, the age range of participants in this study is restricted to young adults (HCP age range 22-37; AOMIC age range 19-26). To assess the generalizability of our results to the whole population, future studies should include subjects with a broader age range. Second, neuroimaging data in general is susceptible to many sources of potential noise (e.g., in-scanner head motion, physiological confound signals, thermal noise) and degrees of freedom in preprocessing strategies can affect study outcomes. However, we have implemented state-of-the art methods and preregistered our analysis strategy to reduce these effects. Third, there are well known limitations to the reconstruction of structural connectivity using fiber tractography that cannot be prevented with our current methodology but need to be considered when interpreting research findings (Schilling et al., 2019; Thomas et al., 2014). Fourth, the observed effect sizes in this study are, according to Cohen (1988), small. However, they lie in the range of what has been recently proposed as maximum reachable effect size for brain-behavior relationships (DeYoung et al., 2022; Marek et al., 2022). Further, note that although cognitive ability measures provide at least moderate reliability (Colom, 2004), the reliabilities of neuroimaging measures are suggested to be rather low (Dennis et al., 2012; Hawco et al., 2018; Noble et al., 2019). Considering that the magnitude of an association between two measures is limited by the multiplication of both measures’ reliabilities and we do not only investigate the association between one behavioral variable and one neurobiological variable, but use a neurobiological variable (SC-FC coupling) already representing a combination of two measures, we consider our observed effect sizes to be close to what can be expected when applying the currently available methodology. Fifth, it is important to note that even though there is evidence that communication models reflect patterns of regional co-activation (Goñi et al., 2014), they only provide putative descriptions of communication processes in the brain. Also, we restricted our analyses to two similarity measures and six communication measures that have been used frequently but various other metrics to assess SC-FC coupling are available as well. Future research could profit from including an extended amount or the combination of communication measures (see Betzel et al., 2022 for a valuable framework allowing to combine multiple models). Further, it is important to highlight that the measure selection process used to generate the NMA masks is not dependent on a threshold, i.e., the measure selected per brain region is not necessarily significantly associated with general cognitive ability which should be considered with respect to our interpretations relating certain communication strategies to GCA. Sixth, FC was estimated using data from resting-state and passive movie watching fMRI. Even though resting-state fMRI has previously been related to individual differences in cognition (Basten et al., 2015), further exploration of the relationship between SC-FC coupling and GCA during active task demands may provide novel information about task-specific communication strategies. Seventh, we limited our analyses to large-scale cortical brain regions as it is not clear so far how different signal-to-noise ratios, as evident in subcortical or hippocampal brain regions (Maugeri et al., 2018; Polders et al., 2011; Vizioli et al., 2021), might affect the computation of communication and similarity measures. However, as subcortical and hippocampal brain regions are suggested to play a critical role in cognitive processing (see e.g., Axmacher et al., 2010; Burgess et al., 2002; Colom et al., 2013), prospective methodological studies comprehensively addressing this question are required.

Overall, we recommend future research to make use of samples with an increased age range, to apply comprehensive replication strategies improving the robustness of research findings, to consider an extended selection of communication measures, to investigate SC-FC coupling during cognitive demands, and to develop means to validly investigate the relationship between SC-FC coupling and GCA also in rather small and medially located brain regions such as the hippocampus.

## 6. Conclusion

This study investigated the question whether individual variations in structural-functional brain network coupling (SC-FC coupling) are associated with individual differences in general cognitive ability (GCA). We used two large openly available datasets and state-of-the-art operationalizations of SC-FC coupling allowing for insights into different neural communication strategies. At the whole-brain level, higher general cognitive ability was linked to stronger SC-FC coupling but only when considering path transitivity as communication strategy. Accounting for region-specific variations in communication strategies within a cross-validated prediction framework enabled significant prediction of individual cognitive ability scores from SC-FC coupling. Finally, despite distinct imaging states, different preprocessing pipelines and another assessment of general cognitive ability, all results replicated in an independent sample and the model developed in the main sample also predicted individual cognitive ability scores in the replication sample. This supports the robustness of our findings as well as the generalizability of the here developed prediction model. Taken together, our results reveal brain region-specific structure-function coupling strategies as neural correlate of individual differences in cognitive ability and provide insights into the basis of efficient information processing as fundamentally implicated in human cognition.

## CRediT authorship contribution statement

**Johanna Popp:** Conceptualization, Methodology, Formal analysis, Writing – Original Draft, Visualization, Funding acquisition. **Jonas Thiele:** Methodology, Writing – Review & Editing. **Joshua Faskowitz:** Methodology, Data curation; Writing – Review & Editing. **Caio Seguin:** Methodology, Writing – Review & Editing. **Olaf Sporns:** Methodology, Writing – Review & Editing. **Kirsten Hilger:** Conceptualization, Methodology, Resources, Writing – Original Draft, Supervision; Funding acquisition.

## Supplementary Material

Supplementary Material associated with this article can be found in a separate document.

## Acknowledgements

The authors thank the Human Connectome Project (Van Essen et al., 2013), WU-Minn Consortium (Principal Investigators: David Van Essen and Kamil Ugurbil; 1U554MH091657) funded by the 16 NIH Institutes and Centers that support the NIH Blueprint for Neuroscience Research, and by the McDonnell Center for Systems Neuroscience at Washington University, for providing data of the main sample, and all contributors to the Amsterdam Open MRI Collection (Snoek et al., 2021) for providing data of the replication samples. We also thank Erhan Genҫ and Christoph Fraenz from the Leibniz Research Centre for Working Environment and Human Factors at the Technical University Dortmund for their contributions to data analysis and development of ideas in the early stages of the project. This research was supported in part by Lilly Endowment, Inc., through its support for the Indiana University Pervasive Technology Institute.

## Abbreviations

AOMIC: Amsterdam Open MRI Collection
BOLD: Blood Oxygen Level Dependent
CoS: Cosine Similarity
FC: Functional Brain Network Connectivity
FG: Flow Graphs
G: Communicability
GCA: General Cognitive Ability
HCP: Human Connectome Project
IST: Intelligence Structure Test
MFPT: Mean First Passage Time
MI: Matching Index
NMA: Node-Measure Assignment
PL: Path Length
PT: Path Transitivity
SC: Structural Brain Network Connectivity
SC-FC Coupling: Structural-Functional Brain Network Coupling
SI: Search Information
TE: Time to Echo
TR: Time to Repetition

