## Supplement for "Structural-Functional Brain Network Coupling Predicts Human Cognitive Ability"

**Supplementary Table S1**

Distribution of coupling measures selected for each brain region in the positive and negative node-measure assignment (NMA) across the seven Yeo networks (Yeo et al., 2011) in the main sample (HCP)

**Positive NMA**

| | CoS | MI | G | MFPT | FG | SI | PT | PL |
|---|---|---|---|---|---|---|---|---|
| VIS | 12 | 0 | 4 | 2 | 9 | 2 | 24 | 4 |
| SOM | 3 | 21 | 3 | 0 | 5 | 0 | 20 | 0 |
| DA | 4 | 19 | 1 | 1 | 11 | 0 | 7 | 1 |
| VA | 0 | 32 | 0 | 0 | 0 | 1 | 16 | 0 |
| LIM | 1 | 1 | 0 | 5 | 6 | 3 | 3 | 5 |
| FP | 0 | 6 | 1 | 7 | 7 | 2 | 15 | 7 |
| DMN | 2 | 7 | 10 | 5 | 20 | 13 | 18 | 12 |
| ALL | 22 (6.15 %) | 86 (24.02 %) | 19 (5.31 %) | 20 (5.57 %) | 58 (16.20 %) | 21 (5.57 %) | 103 (28.77%) | 29 (8.10%) |

**Negative NMA**

| | CoS | MI | G | MFPT | FG | SI | PT | PL |
|---|---|---|---|---|---|---|---|---|
| VIS | 0 | 11 | 4 | 8 | 13 | 13 | 6 | 2 |
| SOM | 1 | 0 | 2 | 12 | 5 | 25 | 5 | 2 |
| DA | 2 | 2 | 6 | 11 | 5 | 5 | 10 | 3 |
| VA | 1 | 0 | 7 | 5 | 11 | 7 | 5 | 13 |
| LIM | 2 | 11 | 5 | 0 | 1 | 1 | 3 | 1 |
| FP | 2 | 10 | 9 | 6 | 5 | 3 | 8 | 2 |
| DMN | 3 | 39 | 6 | 8 | 1 | 8 | 21 | 1 |
| ALL | 11 (3.07 %) | 73 (20.39 %) | 39 (10.89 %) | 50 (13.97 %) | 41 (11.45 %) | 62 (16.20 %) | 58 (16.20 %) | 24 (6.7 %) |

*Note*: Table columns list the number of brain regions in the respective network (table rows) that were assigned to one of the eight coupling measures. VIS = Visual Network; SOM = Somatomotor Network; DA = Dorsal Attention Network; VA = Ventral Attention Network; LIM = Limbic Network; FP = Frontoparietal Network; DMN = Default Mode Network; CoS = Cosine Similarity; MI = Matching Index; G = Communicability; MFPT = Mean First Passage Time; FG

= Flow Graphs; SI = Search Information; PT = Path Transitivity; PL = Path Length; NMA = Node-Measure Assignment.

**Supplementary Table S2**

Prediction performances of 5-fold cross-validated prediction model in the main sample (HCP) when predicting 12 cognitive ability measures that were used to compute the latent *g*-factor

| Test | Instrument | Measure | Prediction Model Performance |
|---|---|---|---|
| 1 | Episodic Memory (Picture Sequence Memory) | PicSeq_Unadj | *r* = .08 (*p* = .020)* |
| 2 | Executive Function/Cognitive Flexibility (Dimensional Change Card Sort) | CardSort_Unadj | *r* = .09 (*p* = .018)* |
| 3 | Executive Function/Inhibition (Flanker Task) | Flanker_Unadj | *r* = .03 (*p* = .173) |
| 4 | Fluid Intelligence (Penn Progressive Matrices) | PMAT24_A_CR | *r* = .17 (*p* = .001)* |
| 5 | Language/Reading Decoding (Oral Reading Recognition) | ReadEng_Unadj | *r* = .17 (*p* < .001)* |
| 6 | Language/Vocabulary Comprehension (Picture Vocabulary) | PicVocab_Unadj | *r* = .19 (*p* < .001)* |
| 7 | Processing Speed (Pattern Completion Processing Speed) | ProcSpeed_Unadj | *r* = .06 (*p* = .054) |
| 8 | Self-regulation/Impulsivity (Delay Discounting) | DDisc_AUC_200 + DDisc_AUC_40K | *r* = .17 (*p* = .001)* |
| 9 | Spatial Orientation (Variable Short Penn Line Orientation Test) | VSPLOT_TC | *r* = .17 (*p* < .001)* |
| 10 | Sustained Attention (Short Penn Continuous Performance Test) | $\frac{SCPT\_TP + SCPT\_TN}{(SCPT\_TP + SCPT\_TN + SCPT\_FP + SCPT\_FN)SCPT\_TPRT}$ | *r* = .05 (*p* = .082) |
| 11 | Verbal Episodic Memory (Penn Word Memory Test) | IWRD_TOT | *r* = .02 (*p* = .305) |
| 12 | Working Memory (List Sorting) | ListSort_Unadj | *r* = .10 (*p* = .016)* |

*Note:* Significance of the prediction model was assessed with a permutation test. *P*-values indicating significant associations are marked with an asterisk (* = *p* < .05).

**Supplementary Table S3**

Relationship between general cognitive ability and brain-average SC-FC coupling in the replication sample (AOMIC)

| **Measure to Compute SC-FC Coupling** | **Replication Sample - AOMIC** *r (p)* |
|---|---|
| Cosine Similarity (CoS) | .05 (.283) |
| Matching Index (MI) | .06 (.179) |
| Communicability (G) | .02 (.712) |
| Mean First Passage Time (MFPT) | .09 (.041) |
| Flow Graphs (FG) | .09 (.028) |
| Search Information (SI) | .05 (.212) |
| Path Transitivity (PT) | .16 (<.001)* |
| Path Length (PL) | .06 (.170) |

*Note:* Replication sample $N = 567$ (AOMIC). The right column lists partial correlations between cognitive ability scores and individual measure-specific brain-average coupling values (measure-specific averages of coupling values from all brain regions) controlled for age, gender, handedness, and in-scanner head motion. Significant associations passing the Bonferroni-corrected threshold (eight comparisons) are marked with an asterisk (* = $p < .006$).

**Supplementary Table S4**

Relationship between general cognitive ability and brain-average SC-FC coupling additionally controlled for total intracranial volume in the main sample (HCP)

| **Measure to Compute SC-FC Coupling** | **Main Sample Not Controlled for Total Intracranial Volume** *r (p)* | **Main Sample Controlled for Total Intracranial Volume** *r (p)* |
|---|---|---|
| Cosine Similarity (CoS) | .05 (.115) | .04 (.200) |
| Matching Index (MI) | .07 (.031) | .08 (.007) |
| Communicability (G) | .04 (.247) | .03 (.331) |
| Mean First Passage Time (MFPT) | .03 (.415) | <.01 (.935) |
| Flow Graphs (FG) | .08 (.010) | .06 (.055) |
| Search Information (SI) | .02 (.588) | .02 (.533) |
| Path Transitivity (PT) | .10 (.002)* | .10 (.002)* |
| Path Length (PL) | .05 (.095) | .04 (.173) |

*Note:* Main sample $N$ = 1030 (HCP). The middle column lists partial correlations between cognitive ability scores and individual measure-specific brain-average coupling values (measure-specific averages of coupling values from all brain regions) controlled for age, gender, handedness, and in-scanner head motion. The right column lists partial correlations between cognitive ability scores and individual measure-specific brain-average coupling values controlled for age, gender, handedness, in-scanner head motion and total intracranial volume. Significant associations passing the Bonferroni-corrected threshold (eight comparisons) are marked with an asterisk (* = $p < .006$).

**Supplementary Table S5**

Post-hoc analysis relating the node-measure assignment (NMAs) masks to the Margulies gradient describing the macroscale cortical organization in terms of uni- and multimodal brain areas

| Node-Measure Assignments (NMAs) | Group-Average of Margulies Gradient (*Mean/N*) | | | | One-Way ANOVA | Post-Hoc Test (Tukey's HSD) | | |
|---|---|---|---|---|---|---|---|---|
| | Diffusion (D) | Path Accessibility (P) | Routing (R) | Similarity (S) | | Group Comparison | Mean Difference | *p* |
| **Positive NMA** | 1.55 (97) | -.16 (124) | 3.37 (29) | -2.20 (108) | *F*(3,354) = 29.26, *p* < .001) | D vs P | -1.70 | .002* |
| | | | | | | D vs R | 1.82 | .068 |
| | | | | | | D vs S | 3.75 | < .001* |
| | | | | | | P vs R | 3.53 | < .001* |
| | | | | | | P vs S | 2.04 | < .001* |
| | | | | | | R vs S | 5.57 | < .001* |
| **Negative NMA** | -1.18 (130) | -.65 (120) | -2.49 (24) | 3.36 (84) | *F*(3,354) = 37.88, *p* < .001). | D vs P | .53 | .613 |
| | | | | | | D vs R | - 1.31 | .311 |
| | | | | | | D vs S | - 4.54 | < .001* |
| | | | | | | P vs R | - 1.84 | .076 |
| | | | | | | P vs S | - 4.01 | < .001* |
| | | | | | | R vs S | - 5.85 | < .001* |

*Note:* Values from the Margulies gradient associated with brain regions that were assigned to each of the four coupling measure groups (diffusion, path accessibility, routing, and similarity) were compared across groups using a one-way ANOVA. Column 1-5: A negative group mean indicates higher prevalence of brain regions assigned to this measure group (D, P, R, S) in unimodal areas, while a positive group mean specifies higher prevalence in multimodal areas. The table lists mean values of the Margulies gradient for each coupling measure group, number of brain regions assigned to each group (*N*), results of the one-way ANOVA and of the post-hoc test (Tukey's HSD). Significant group differences are marked with an asterisk (* = *p* < .05). NMA = Node-Measure Assignment; D = Diffusion; P = Path Accessibility; R = Routing; S = Similarity.

**Supplementary Table S6**

Prediction performances of the 5-fold cross-validated prediction model in the main sample (HCP) when one coupling measure is excluded from the selection process at a time

| Test | Excluded Measure | Prediction Model Performance |
|---|---|---|
| **Baseline** | None | $r$ = .25 ($p$ < .001)* |
| **1** | Path Length (PL) | $r$ = .25 ($p$ < .001)* |
| **2** | Communicability (G) | $r$ = .25 ($p$ < .001)* |
| **3** | Cosine Similarity (CoS) | $r$ = .25 ($p$ < .001)* |
| **4** | Search Information (SI) | $r$ = .25 ($p$ < .001)* |
| **5** | Path Transitivity (PT) | $r$ = .26 ($p$ < .001)* |
| **6** | Matching Index (MI) | $r$ = .21 ($p$ < .001)* |
| **7** | Mean First Passage Time (MFPT) | $r$ = .26 ($p$ < .001)* |
| **8** | Flow Graphs (FG) | $r$ = .25 ($p$ < .001)* |

*Note:* Significance of the prediction model was assessed with a permutation test. *P*-values indicating significant associations are marked with an asterisk (* = $p$ < .05).

**Supplementary Table S7**

Prediction performances of the 5-fold cross-validated prediction model in the main sample (HCP) when one prediction feature is excluded from the model at a time

| Test | Features Used | Prediction Model Performance |
|---|---|---|
| **Baseline** | Positive and Negative NMA | $r$ = .25 ($p$ < .001)* |
| **1** | Positive NMA only | $r$ = .21 ($p$ = .001)* |
| **2** | Negative NMA only | $r$ = .03 ($p$ = .238)* |

*Note:* Significance of the prediction model was assessed with a permutation test. *P*-values indicating significant associations are marked with an asterisk (* = $p$ < .05).

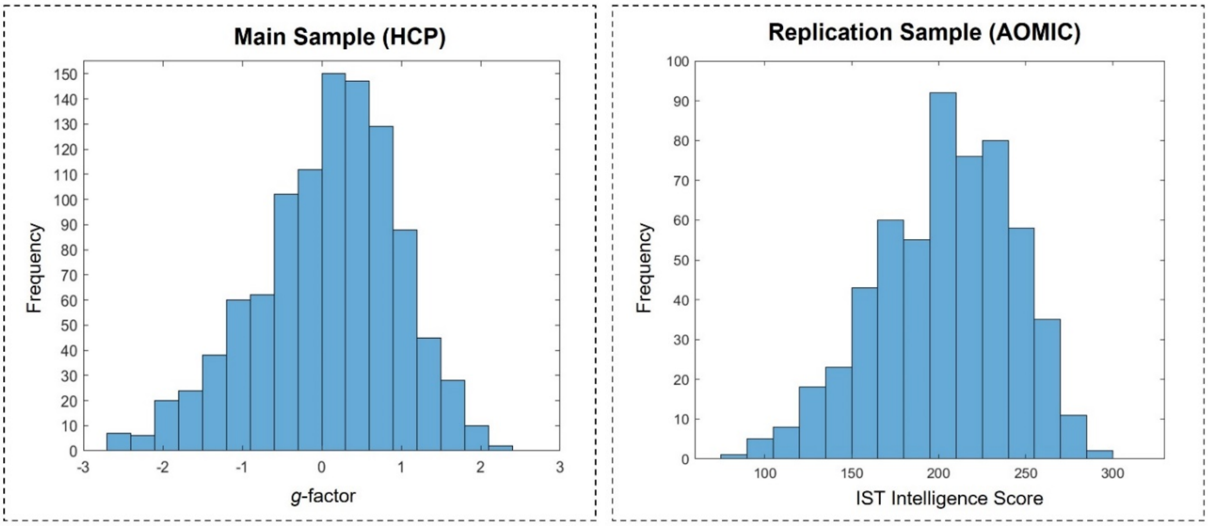


**Supplementary Fig. S1.** Distribution of cognitive ability scores in the main sample (*N* = 1030, HCP, left) and in the replication sample (*N* = 567, AOMIC, right). Cognitive ability in the main sample was operationalized as latent *g*-factor (see Methods). The *g*-factor ranged between -2.60 and 2.40 (*M* = 0.08; *SD* = 0.89). In the replication sample, cognitive ability was assessed using sum scores of the Intelligence Structure Test (IST, Beauducel et al., 2010), which ranged between 78 and 295 (*M* = 202.85; *SD* = 39.01). HCP = Human Connectome Project; AOMIC = Amsterdam Open MRI Collection; IST = Intelligence Structure Test.

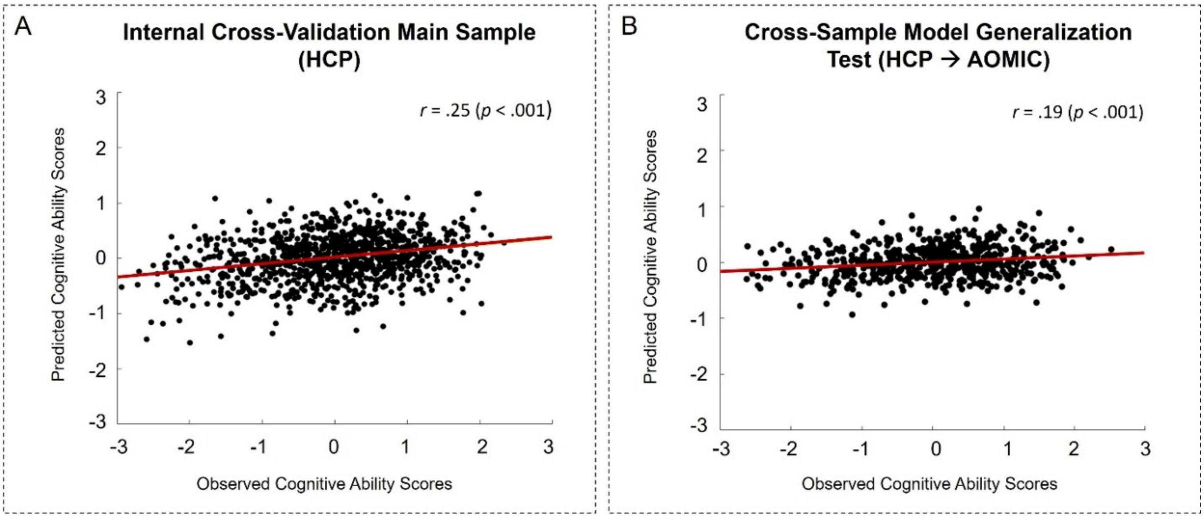


**Supplementary Fig. S2.** Scatterplots illustrating the relationship between predicted and observed cognitive ability scores of the cross-validated prediction model applied within the main sample (*N* = 1030, HCP) and for the cross-sample model generalization test (*N* = 567, AOMIC). (A) Significant association between predicted and observed cognitive ability scores from a 5-fold internally cross-validated prediction model using mean coupling values from positive and negative node-measure assignment (NMA) as predictors in the main sample: $r = .25$, $R^2 = .06$, $p < .001$ by permutation test. (B) Significant association between predicted and observed cognitive ability scores of the cross-sample model generalization test, i.e., the prediction model built in the main sample (HCP) was used to predict cognitive ability scores in the replication sample (AOMIC): $r = .19$, $R^2 = .04$, $p < .001$ by permutation test. Lines of best linear fit are plotted in red. HCP = Human Connectome Project; AOMIC = Amsterdam Open MRI Collection.

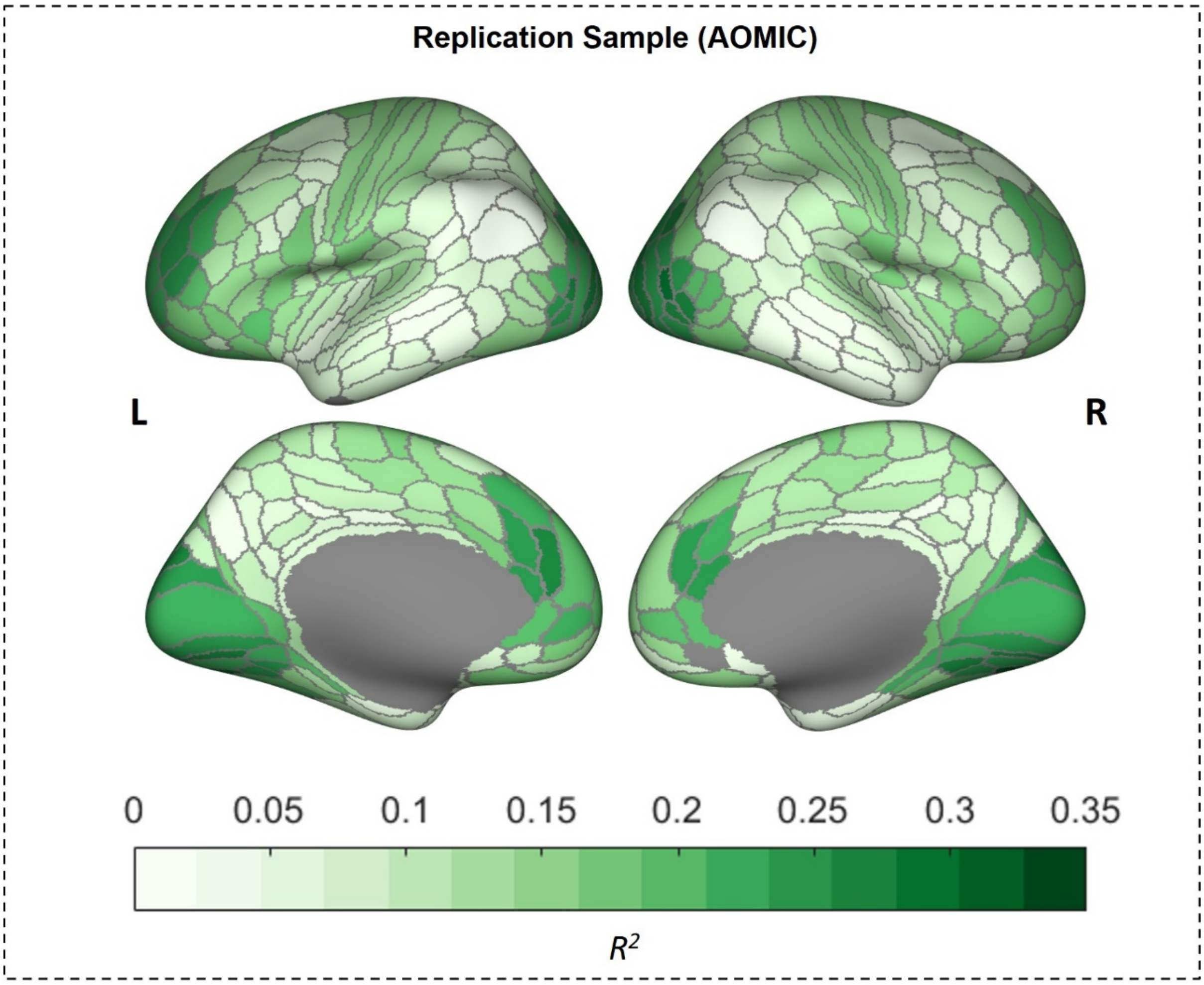


**Supplementary Fig. S3.** Group-average whole-brain pattern of SC-FC coupling strength. This figure illustrates the cortical distribution of coupling strength based on the region-specific similarity or communication measure (computed based on structural connectivity) able to explain the highest amount of variance in functional connectivity most frequently across all participants in the replication sample (AOMIC). This group-general mask was subsequently used to extract individual region-specific coupling values ($R^2$) and a group-average map of regional coupling strength was created by averaging across all participants' coupling values. Note that this approach was solely used for visualization purposes but not for further statistical analyses. In correlative and predictive analyses, the coupling measure was determined by a group-general mask (i.e., the positive and negative NMAs) based on the strongest association with GCA. These latter masks were also cross-validated in all predictive approaches to prevent any data leakage between training and test samples.

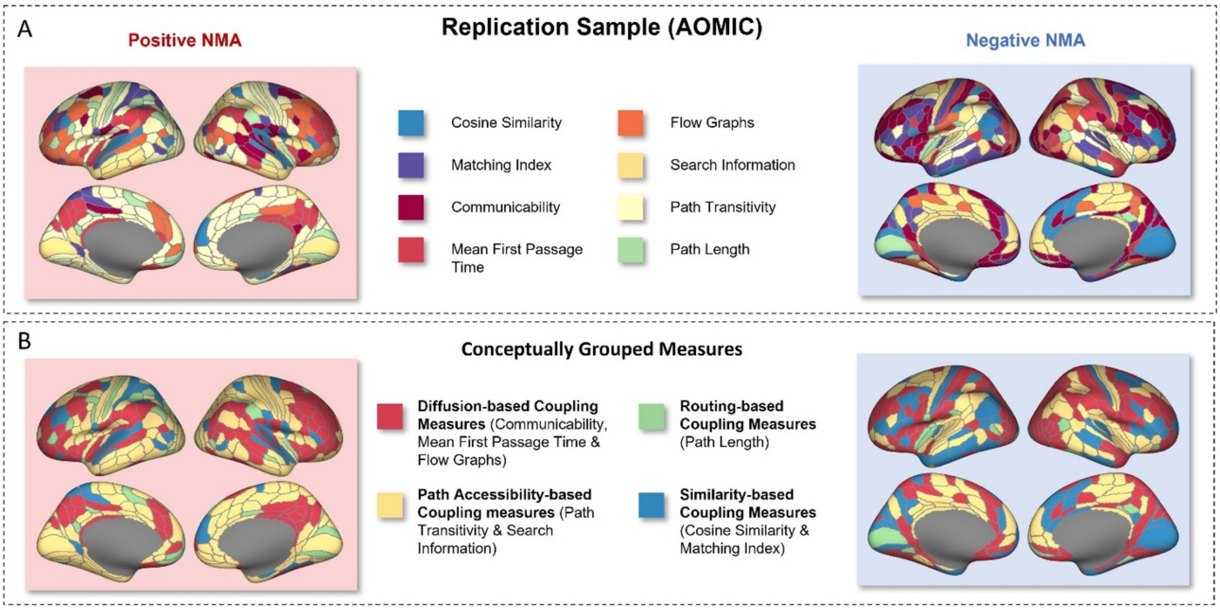


**Supplementary Fig. S4.** General cognitive ability is associated with brain region-specific SC-FC coupling also in the replication sample (AOMIC). (A) Group-based positive and negative node-measure assignment (NMA) masks built in the replication sample. These masks were created by identifying the coupling measure with the largest positive and negative magnitude association with GCA ($r_G$) per brain region across all participants of the replication sample (*N* = 567), depicting the measure per brain region best explaining individual differences in GCA. Note that to thoroughly prevent overfitting, for the 5-fold internal cross-validation, NMA masks were created separately for each training sample of each cross-validation loop and could thus slightly differ from the masks shown in this illustration. Individual-specific coupling values ($r_C$) were then extracted with these masks, i.e., the group-based NMA mask defined which individual-specific coupling value (one out of eight different coupling values, see Methods) was extracted for each brain region and used for further analyses. (B) For illustration purposes, coupling measures were grouped based on conceptual similarity of the proposed signaling mechanism. The grouped positive NMA mask revealed that coupling measures based on diffusion were chosen predominantly in frontal regions and coupling measures based on path accessibility were selected for a widely distributed set of brain regions. The grouped negative NMA mask showed that coupling measures based on routing were only very seldomly selected, while coupling measures based on diffusion were chosen for many brain regions. AOMIC = Amsterdam Open MRI Collection, NMA = Node-Measure Assignment.

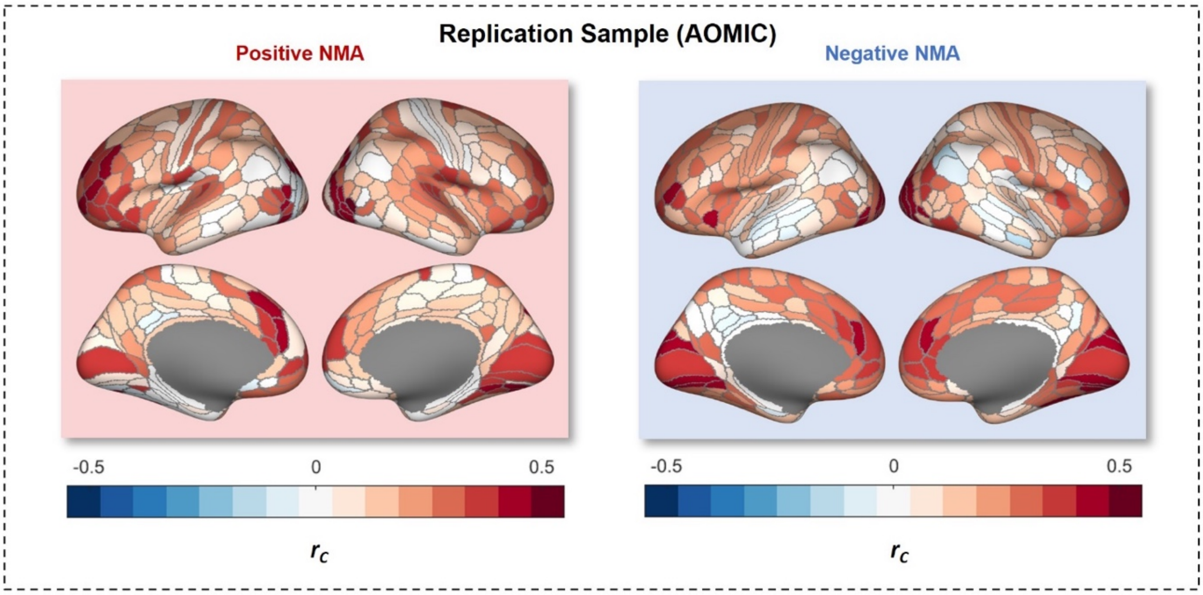


**Supplementary Fig. S5.** Group-average region-specific SC-FC brain network coupling strength corresponding to the node-measure assignment (NMA) masks built in the replication sample (AOMIC, $N$ = 567). These masks were created by identifying the coupling measure with the largest positive and negative magnitude associations with GCA ($r_G$) per brain region across all participants of the complete replication sample. For each participant, individual-specific coupling values were extracted using the group-based NMA masks, resulting in two individual brain maps (for the positive and negative NMA mask, respectively) containing one coupling value ($r_C$) for each brain region. This figure illustrates a group-average map of these individual regional coupling values. Note however, that these values were not used in further analyses. AOMIC = Amsterdam Open MRI Collection; NMA = Node-Measure Assignment.

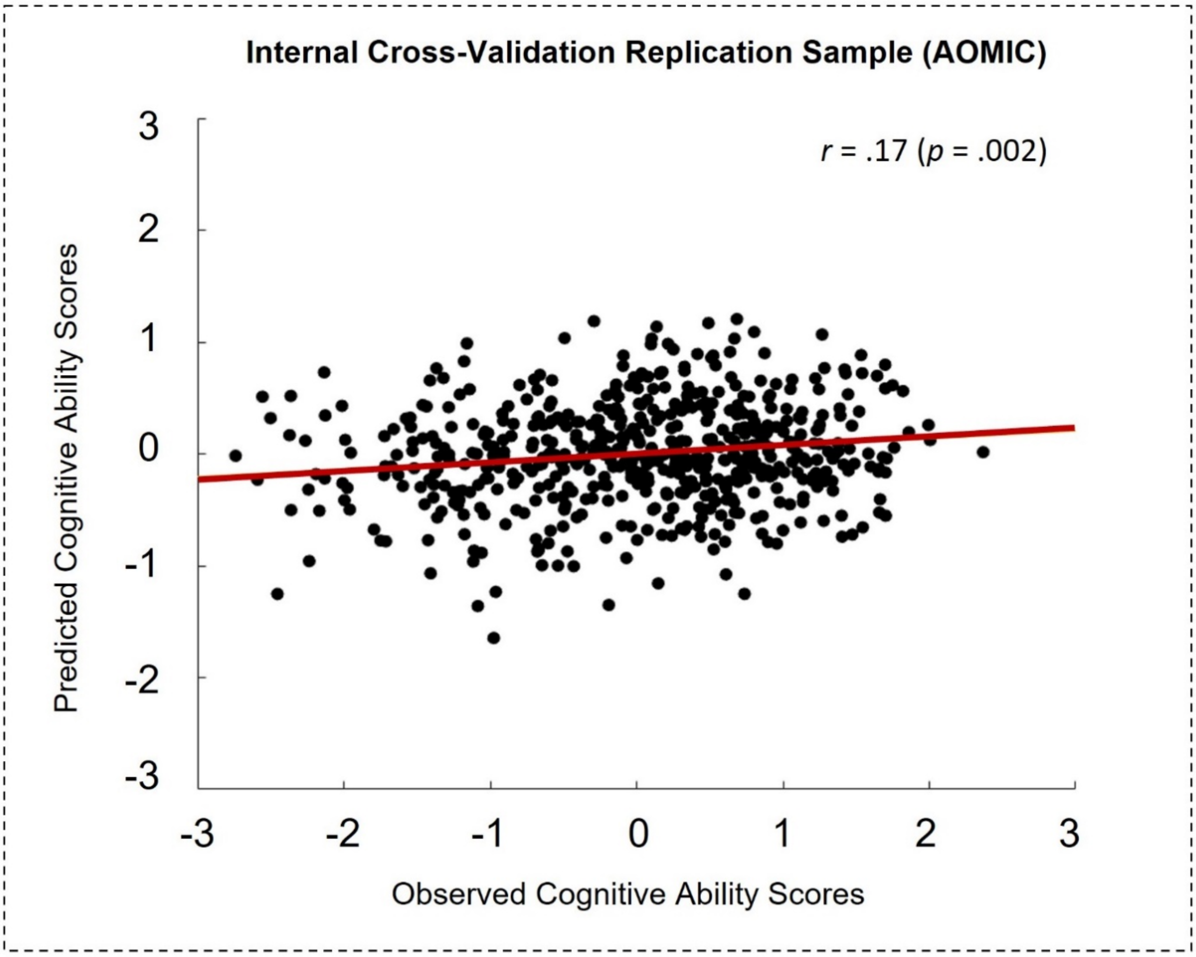


**Supplementary Fig. S6.** Scatterplot illustrating the association between observed and predicted cognitive ability scores in the cross-validated prediction model within the replication sample (AOMIC; $N$ = 567). Significant association between predicted and observed cognitive ability scores from a 5-fold internally cross-validated prediction model using individual-specific mean coupling values from positive and negative node-measure assignment (NMA) as predictors within the replication sample: $r$ = .17, $R^2$ = .03, $p$ = .002 by permutation test. Lines of best linear fit are plotted in red. AOMIC = Amsterdam Open MRI Collection.

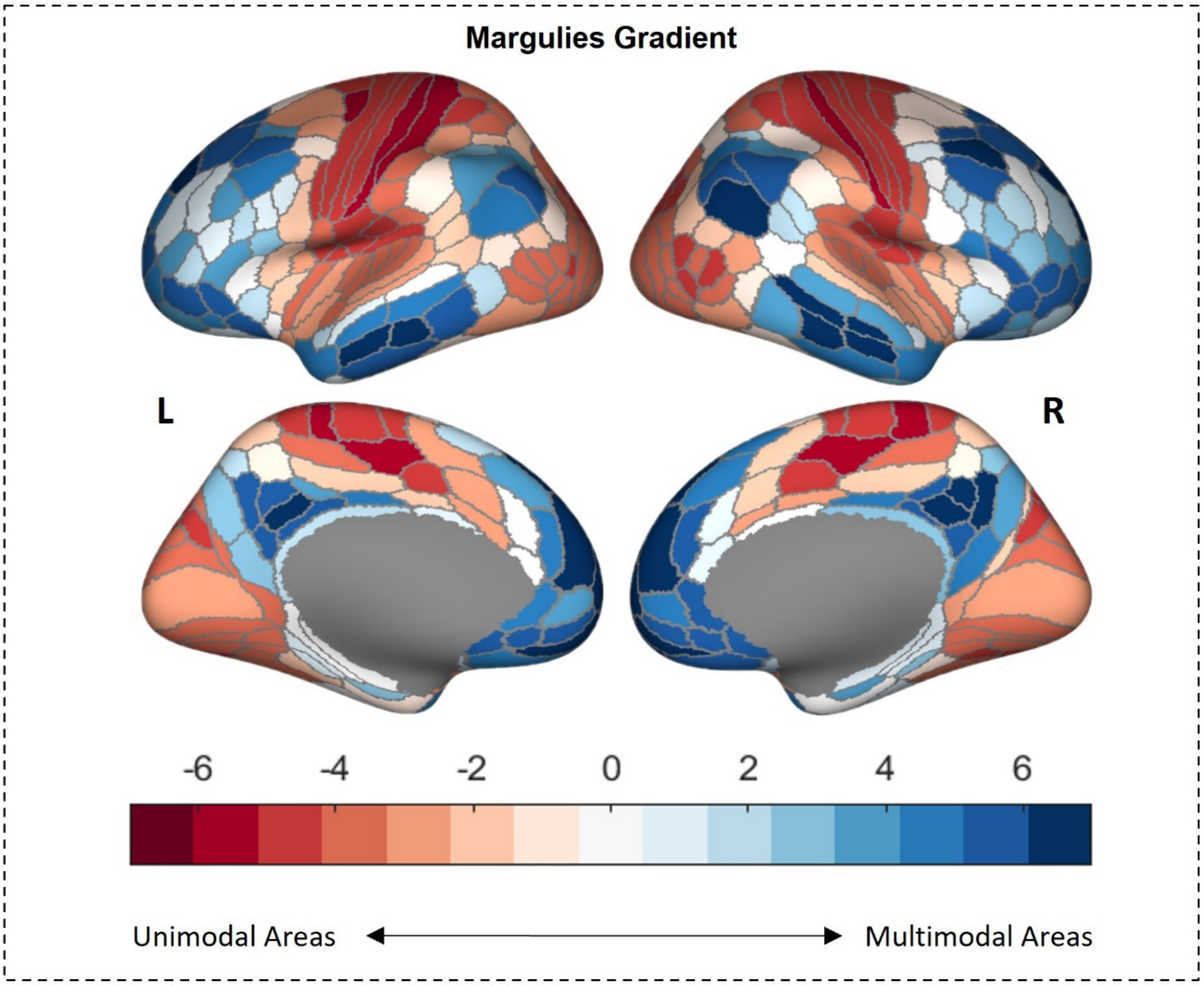


**Supplementary Fig. S7.** Visual representation of the Margulies gradient. The gradient (Margulies et al., 2016) describes the macroscale cortical organization and situates each brain region on a continuum between unimodal regions serving primary sensory and motor tasks (negative values, red) and multimodal regions serving complex heteromodal processing (positive values, blue).

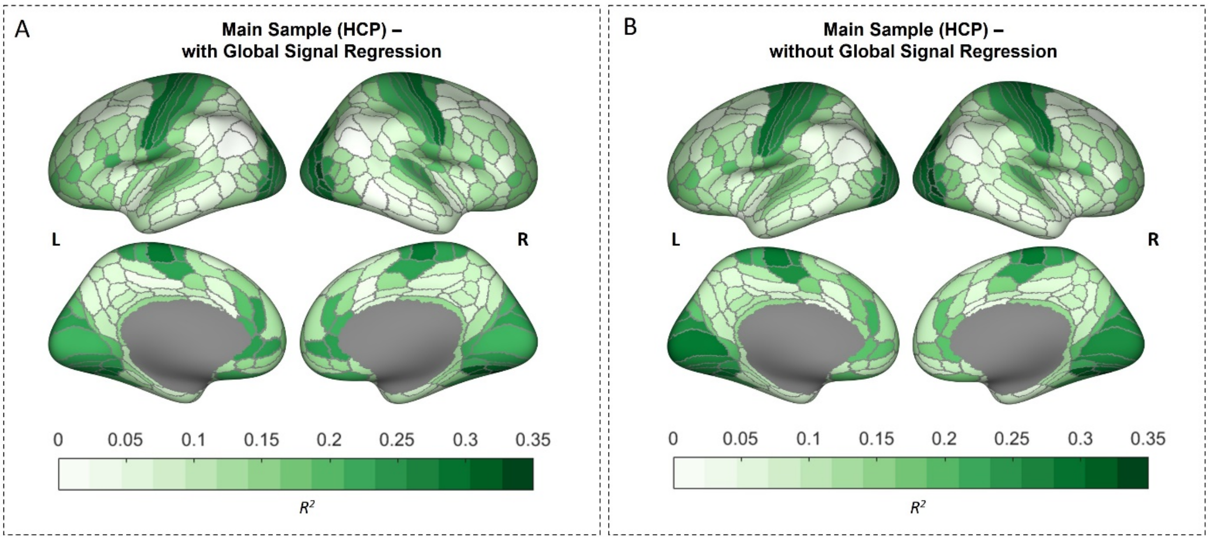


**Supplementary Fig. S8.** Group-average whole-brain pattern of SC-FC coupling strength with and without global signal regression in the fMRI preprocessing strategy. (A) Whole-brain pattern of coupling strength with inclusion of global signal regression in the preprocessing strategy. (B) Whole-brain pattern of coupling strength without inclusion of global signal regression in the preprocessing strategy.

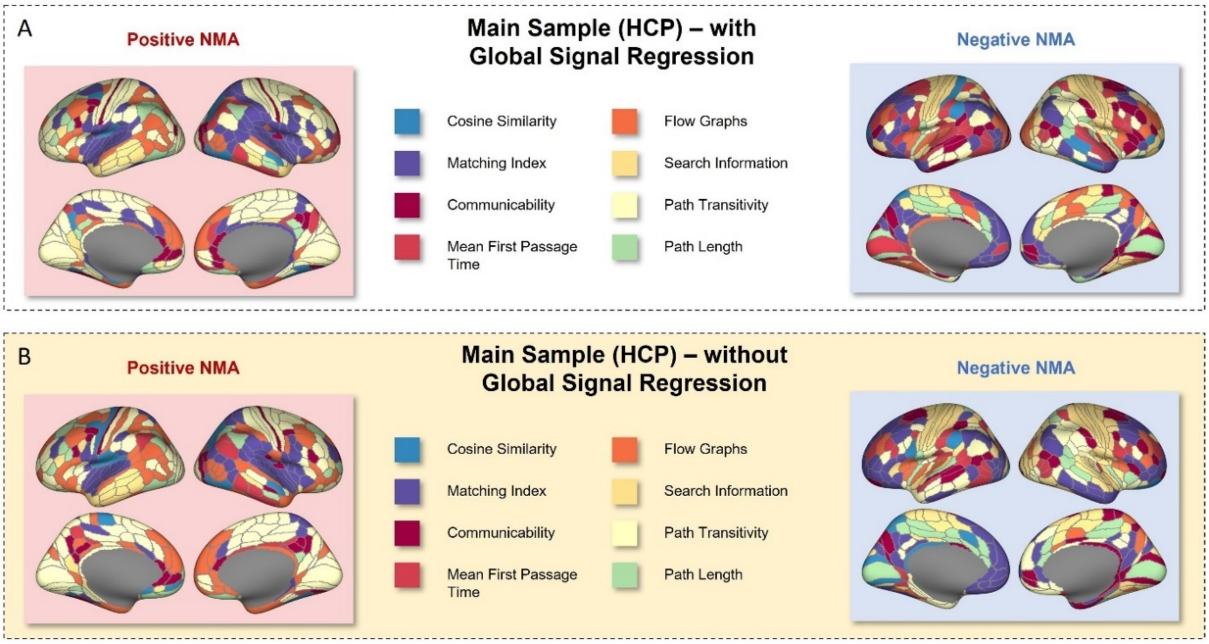


**Supplementary Fig. S9.** Control analysis comparing group-based positive and negative node-measure assignment (NMA) masks that were computed with and without the implementation of global signal regression in the preprocessing strategy of the fMRI data. (A) Positive and negative NMAs computed from SC-FC coupling data including global signal regression in the preprocessing strategy. (B) Positive and negative NMAs computed from SC-FC coupling data without global signal regression in the preprocessing strategy.
